# Mass flowering and flowering asynchrony characterise a seasonal herbaceous community in the Western Ghats

**DOI:** 10.1101/2024.11.10.622833

**Authors:** Saket Shrotri, Sukhraj Kaur, Rahul Dhargalkar, PV Najla, Vinita Gowda

## Abstract

Flowering synchrony within a community may be facilitated by climatic factors and by ecological interactions that promote shared pollination services. In contrast, flowering asynchrony is promoted when there is inter-species competition for pollinators. In this multiyear study, we analyse the flowering phenology of a seasonal, herbaceous community (Kaas plateau) in Western Ghats, India to identify environmental constraints that may influence flowering synchrony. We addressed the questions: (a) Is flowering seasonality correlated with climatic factors? (b) Is there evidence for flowering synchrony within the community? (c) Do plant-pollinator interactions shift with flowering phenology?

In Kaas, we recorded flowering phenology of 76 herbaceous species and found that climatic factors influenced their flowering phenology. We also identified the community to be composed of a few mass flowering (MF) species (∼30%) and several non-mass flowering (nMF) species (∼70%). Using two novel synchrony indices, temporal overlap (SI_temp_) and synchrony in abundance (SI_abd_), we also identified higher asynchronous flowering within the community than expected. Notably, species sharing the same floral colour, showed marked absence of synchrony, thus suggesting that competition and not pollinator-mediated facilitation drives flowering asynchrony within Kaas. Finally, pollination networks were observed to shift with flowering abundances within the community.

Our findings reveal that even seasonal landscapes like the laterite plateaus, despite their short flowering season that last only 4-5 months, exhibit an overall asynchronous flowering phenology. And, the synchronous flowering for which the Kaas plateau is famous, was noted to be mostly due to a few mass flowering species that alter across years.

## 1. INTRODUCTION

Flowering phenology i.e., the timing and pattern of flower production plays a pivotal role in the reproductive success of plants. Among individuals of the same species, selection often favours synchrony in flowering to maximize pollinator attraction (Gentry, 1974; Bawa et al., 1985; Rathcke and Lacey, 1985; Kudo, 2006; Elzinga et al., 2007) and facilitate cross-pollination while minimizing heterospecific pollen transfer (HPT; Ashton, 1969; Augspurger, 1983; Ashton et al., 1988; Sakai et al., 1999). However, at the community level, flowering patterns may be shaped by contrasting evolutionary pressures. For instance, flowering synchrony among co-occurring species may emerge due to common environmental triggers (e.g., seasonal rainfall) or due to adaptive strategies that amplify floral displays, that can enhance pollinator attraction through facilitation (Parrish and Bazzaz, 1979; Ims, 1990; Sakai et al., 1999; Nagahama and Yahara, 2019) or satiate predators in masting (Janzen, 1971; Zwolak et al., 2022). On the contrary, flowering asynchrony (or staggered flowering) may be favored in communities with high niche overlap, as it reduces competition for pollinators (Schemske, 1981; Aizen and Vázquez, 2006; Lasky et al., 2016). In tropical communities, the interplay between these two opposing forces—facilitation versus competition—is essential to understand and predict phenological patterns that shape plant communities.

Mass flowering (MF) is an extreme example of synchronous flowering and is typically viewed as a community-wide phenomenon where multiple species flower simultaneously in a supra-annual manner (Ashton et al., 1988; Appanah, 1993). However, since these community-wide MF events arise from collective contributions of flowering phenologies of individual species, the term MF has also been used to refer to species-specific events. That is, when a single species shows hyper-abundant flowering in a landscape resulting in a localized high-density flowering (i.e., mass flowering) event (Baker, 1959; Gentry, 1974; Augspurger, 1983). To the best of our knowledge, there are no studies that unify the concepts between MF observed within a species and MF observed within a landscape. Such unification is especially critical to understand dynamics of communities predominantly composed of annuals, and thus there is a strong need to establish the species-specific definition for MF.

One method to unify the above two definitions of MF is to first define a species-specific MF event and then use it to explain the community-wide or landscape-scale MF phenomenon. In this study, a species is identified as MF when it displays high flowering density relative to all species within the community (Box 1). This density-dependent definition acknowledges that a species may display inter-annual variation in its flowering, and thus it may or may not display MF in consecutive years. By clarifying this subtle yet critical distinction, we emphasize that MF is not synonymous with masting (which emphasizes interannual variability in reproductive investment) but rather a measurable trait that contributes to the overall phenological pattern of the community or a species (Box 1).

Thus, in a community with a few MF species, nMF species may benefit by co-flowering with MF species and therefore a higher synchrony may be predicted between MF and nMF species resulting in a co-ordinated, amplified floral display (Tepedino and Stanton, 1981; Augspurger, 1983; Gumbert et al., 1999; Sakai, 2002; Bergamo et al., 2020). Within a community, synchrony in flower production among MF species may result in convergence of floral traits, particularly in floral colour. When species with similar floral hues synchronise their flowering, their visual signals to pollinators can be amplified. For instance, MF species may facilitate pollination in non-mass flowering (nMF; low flowering density) species through trait matching, as per the pollinator-mediated facilitation hypothesis (Bergamo et al., 2020). Thus, co-flowering species with shared floral colours may show higher flowering synchrony to optimise pollinator attraction and the resulting plant-pollinator interactions may also show temporal variation with flowering peaks.

We test the above predictions of synchrony in flowering phenology in a seasonally flowering, high-altitude laterite plateau (‘sky island’) from the Western Ghats, India. The laterite plateaus host a unique set of annual herbaceous taxa with exceptional endemism (Lekhak and Yadav, 2012) and conspicuous mass-flowering events (Figure 1). We used phenology transects to track flowering of 76 herbaceous species (∼90% of plateau’s herbaceous flora) over three years and identify MF and nMF species. To test the role of climatic factors and the role of competition and facilitation in floral colour-mediated synchrony between MF and nMF species, and corresponding shifts in plant-pollinator interactions, we addressed the following questions:

a. Do climatic factors explain community-wide flowering seasonality?
b. Do MF species synchronize with each other or with nMF species?
c. Do convergent floral color patterns operate at a community-level, resulting in distinct synchrony among co-flowering species based on their floral color?
d. How do plant-pollinator networks shift temporally in response to flowering phenology?

**Figure 1:**
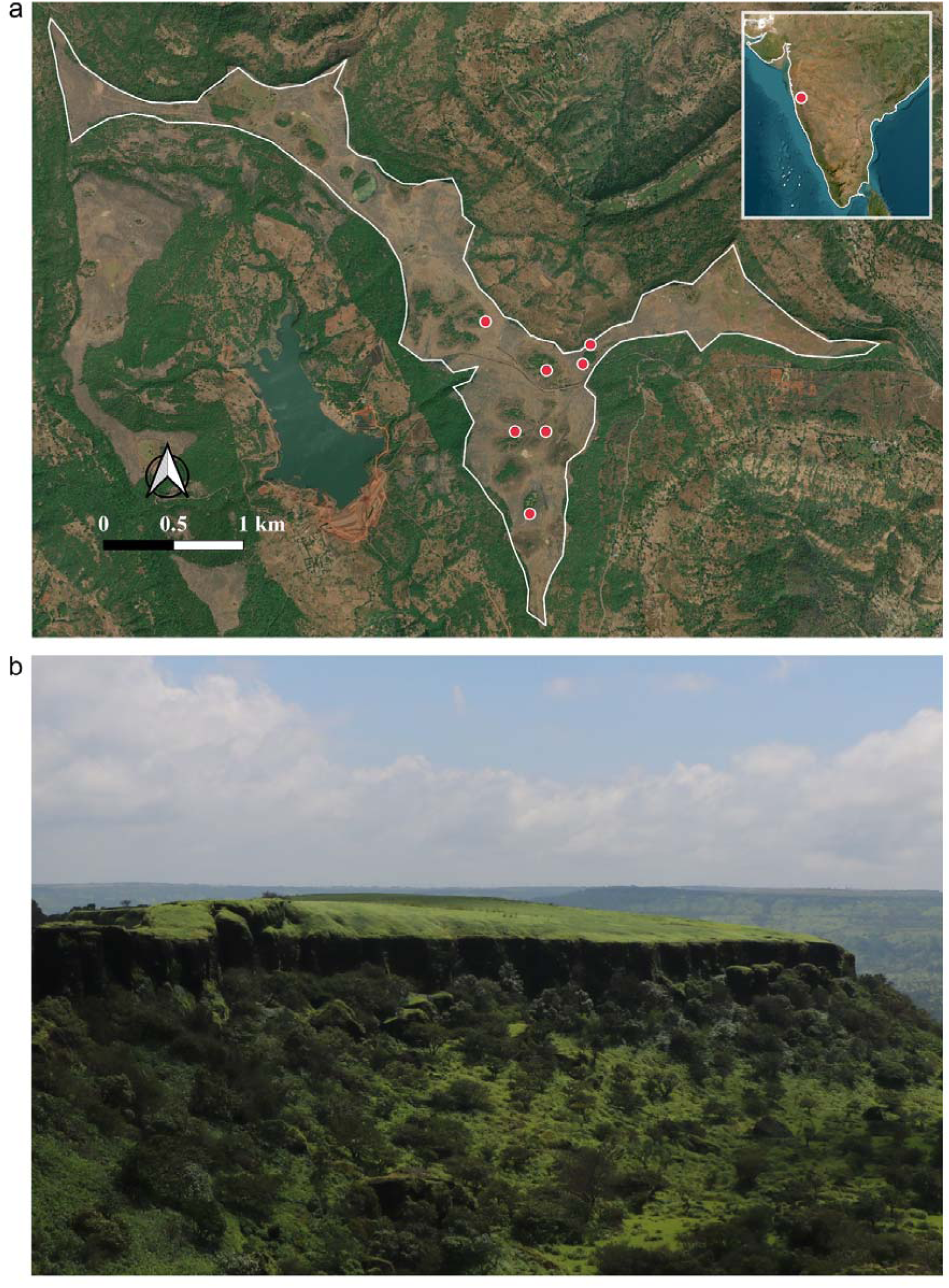
Study site and habitat: (a) Map demarcating the boundary of Kaas plateau, with the locations of phenology transects marked in red (*n* = 7). The inset map displays peninsular India, highlighting the geographical location of the study site (red circle). (b) A typical sky island habitat of Northern Western Ghats, Maharashtra, India, with distinct geographical isolation from the surrounding vegetation.

We predict that the onset of Indian monsoon may impose a stringent environmental filter resulting in flowering synchrony within the community. We also predict that facilitation may promote floral colour-mediated synchrony between MF and nMF species and that shifts in flowering phenology will be accompanied by corresponding shifts in plant–pollinator interactions.

## 2. METHODS

### 2.1. Study area: geography and climate

We conducted all phenological and pollinator observation studies at Kaas plateau (कास पठार, 17°43’12.4” N, 73°49’28.1’’ E; Figure 1a) between 2018 and 2020. Kaas is a high-altitude lateritic plateau situated at 1250 m a.s.l. in the Northern Western Ghats, India (Lekhak and Yadav, 2012). We identify the laterite plateaus as ‘sky islands’ owing to their prominent cliff edges, steep slopes, and isolated biotic communities separated by surrounding valleys (Figure 1b). Sky islands of the Western Ghats are characterised by rocky outcrops with minimal soil substrate (Watve, 2013). They experience extreme environmental conditions such as a prolonged dry season lasting seven months from November to May, followed by a wet season from June to October, which marks the Indian monsoon. Hence, these plateaus primarily support herbaceous plant species, many of which are endemic and exhibit flowering during the monsoonal or post-monsoonal period (Figure S1).

### 2.2. Climatic parameters

Due to the absence of local weather stations within 100 km of the study site, climatic data were retrieved from the NASA POWER database (https://power.larc.nasa.gov) for January 2018–December 2020 with ½° latitude, ⅝° longitude resolution. We obtained daily variations in five climatic parameters: Rainfall (mm), maximum (T_max_), minimum (T_min_) and average (T_avg_) temperature at two metres (°C), relative humidity at two metres (%), and surface soil wetness (proportion). To account for delayed climatic effects on flowering phenology, we employed Generalized Additive Models for Location, Scale, and Shape (GAMLSS; Rigby and Stasinopoulos, 2005) with cubic spline smoothing and forward stepwise selection (GAIC criterion). Each model assumed a Negative Binomial Type I (NBI) distribution to accommodate overdispersed floral count data with excess zeros and the RS algorithm (Stasinopoulos and Rigby, 2008) was used to obtain the estimates of the climatic predictors.

Five separate models were constructed to evaluate the independent effects of rainfall, maximum/minimum temperatures, average temperature, relative humidity, and soil wetness. Lagged climatic predictors (optimum lags were determined via cross-correlation analysis) were incorporated to assess delayed impacts. To mitigate multicollinearity among parameters, each climatic variable was analyzed independently. Climatic data were temporally aligned with floral census intervals (10-day resolution) prior to analysis. All analyses were conducted in R v4.4.1 (R core team, 2022) and the codes are available via GitHub repository (see open research statement).

### 2.3. Flowering phenology

We conducted periodic flowering censuses in each study year between August to December at 15-day intervals in 2018 and between June to November at 10-day intervals in 2019 and 2020 using seven transects (20 x 2 m^2^; Figure 1a) that resulted in a total sampled area of 280 m^2^ per census. In addition to this, in 2015, 2016 and 2017, the flowering phenology of several species (MF and nMF) were documented in the study site using non-systematic surveys. We quantified the flowering abundance of all herbaceous taxa by counting the total number of flowering individuals present within the transect, and we use this term interchangeably with flowering density (used mostly for a species than for community phenomenon) as it is also measured within a fixed area. We identified a species to display ‘high flower density’ if its flowering density exceeded 1.64 times the standard deviations above the community-wide mean flowering density (i.e., z-score of 1.64, ensuring that any value above this threshold exceeds 95th percentile; Zar, 1999). This threshold is temporally dynamic in nature as it is dependent on the total flowering abundance of the entire community, and captures real-time shifts in flowering. Species that did not meet this threshold were classified as species that displayed a ‘low flower density’. The high and low flower density thresholds were quantified for each species within the transect during each census across years (Box 1; Table S1). This classification recognizes mass flowering as a high flower density event, irrespective of whether it occurs in a temporally short, or long flowering period [Gentry’s type 3 and 4 (1974)] and aligns with the classical definition of mass flowering (Box 1).

To test whether species may be synchronised by their floral colour, we categorised all recorded species into three groups based on their floral spectral reflectance (Figure S2). While, these groups are named after the colours perceived by human vision, such as Whites (WH), Pinks and Purples (PK-PL), and Yellows (YL), the three colour categories were defined using cumulative spectral data from a total of 13 species (Figure S2); although we did not perform spectral analyses for all the species. The spectral reflectance values for Whites (WH) range from approximately 400 to 700 nm with high reflectance across the spectrum, Pinks and Purples (PK-PL) show distinct peaks in the 400-500 nm and 600-700 nm ranges, and Yellows (YL) exhibit high reflectance in the 500-600 nm range (Figure S2).

### 2.4. Flowering synchrony

We quantified flowering synchrony using the phenological data collected in 2020. Due to limitations of known synchrony indices (Box 2), we propose two new synchrony indices to measure temporal synchrony (SI_temp_: flowering overlap) and synchrony in abundance (SI_abd_ : synchrony of flowering intensity) between two species (see Box 2 for a detailed explanation). Synchrony in flowering time (SI_temp_), is modified from Primack (1980) and quantifies flowering synchrony among species and is defined as the ratio of the flowering overlap between species 1 and species 2 with the total flowering duration of the two species (equation 1; Box 2).

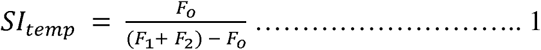

where, *F_1_* and *F_2_* represent the total flowering duration (number of days in flowering) for species 1 and species 2, respectively, and *F_o_* represents the temporal overlap (number of days when two species co-flower) in flowering. SI_temp_ ranges from 0 to 1, where zero represents no temporal overlap, while 1 represents a complete overlap of flowering duration between two species (i.e., when F_o_ = F_1_ = F_2_; Box 2). We quantified SI_temp_ between all species-pairs recorded in the phenology census.

Whenever two species showed temporal overlap (i.e., SI_temp_ > 0 or *F_o_* ≠ 0), we calculated the synchrony in flowering abundance (SI_abd_) between them. This metric provides an index for the relative flowering abundance of two co-flowering species. To measure SI_abd_ between two species, we calculated the ratio between the less abundant species and the more abundant species at any given census. This was summed across all censuses for the species pair (n; equation 2; Box 2).

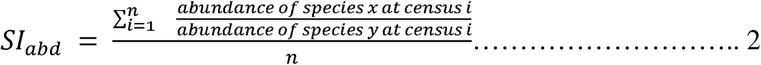

The value of SI_abd_ ranges from 0 to 1, where one represents an equal flowering abundance of the two species when they co-flower. For both indices, synchrony values below 0.4 were considered asynchronous, while values above 0.6 were considered synchronous, and values between 0.4 and 0.6 were characterised as indeterminable (Oleques et al., 2017). Our proposed metric for SI_temp_ and SI_abd_ were also tested in an independent study by Oleques et al., (2017) where the data was appropriate.

We analysed flowering synchrony: (a) among all species in the community (b) among MF and nMF species pairs, using data from 2018, 2019, and 2020; and we used observations from 2020 to calculate synchrony (c) within and among the three floral colour groups (WH, PK-PL, and YL). Synchrony indices were calculated and analysed using R v4.4.1 and the codes are available via GitHub repository (see open research statement).

### 2.5. Plant-pollinator network

We conducted manual pollinator observations on a total of 46 flowering species (∼ 52% of total species recorded in the phenology transects) between August 2018 and October 2019. Observations were carried out between 0600 hrs to 1600 hrs using a total of 339 observation slots of 10–20-minutes. A floral visitor was identified as a pollinator only when it came in contact with the reproductive structures of the flower. We recorded the taxonomic identity of the visiting pollinator type, the total number of visits in a single visitation bout, and the number of flowers in the observation plot. To characterise broad plant-pollinator patterns within the community we grouped pollinators into seven categories: bees, flies, beetles, butterflies, wasps, moths, and ants.

To capture temporal shifts in plant-pollinator interactions during the flowering season, we divided the flowering season into three sub-seasons based on floral abundance and diversity: (a) pre-peak (August to mid-September) with low abundance and diversity, (b) peak (mid-September to early October) with the highest abundance and diversity, and (c) post-peak (mid-October to November) with moderate to low abundance and diversity. We then calculated pollinator visitation rates (number of visits per flower per hour) for every plant-pollinator pair within each sub-season. The visitation rates were used to construct interaction matrices, with plant species as rows (lower trophic level) and pollinators as columns (higher trophic level). We constructed three sub-seasonal networks using the *bipartite* package (Dormann et al., 2009) in R (version 4.4.1) and created a single cumulative network for the entire flowering season to understand the community-wide nature of plant-pollinator interactions. For all networks, we calculated the following indices: links per species (LPS), Shannon’s diversity index (H’), degree of specialisation (H_2_’), and network modularity. We also computed the mean number of shared partners and niche overlap among plant species, as well as species strength and specialisation index (d’) for pollinator types. Finally, we tested the significance of the network indices by comparing the observed network values to a null model (Vázquez and Aizen, 2003) using 10,000 iterations with a 95% confidence interval.

## 3. RESULTS

### 3.1. Environmental seasonality and effect of climatic factors

Based on the climatic and environmental features, we identified three distinct seasons: cold-dry (November to February), hot-dry (March to May), and the wet season (June to October, which coincides with the Indian monsoon; Figure 2a). During the three-year study period (2018 to 2020), daily temperatures fluctuated from 9.4 to 38.7 °C in the cold-dry season and 14 to 44 °C in the hot-dry season. Temperature fluctuations were lowest during the wet season (ranging from 14.6 to 32.8 °C), compared to other seasons. Peak rainfall was recorded in July and August, with an average annual precipitation of ∼2500 mm (Figure 2a). GAMLSS analysis showed that flowering phenology could be predicted by rainfall, temperature, humidity and soil wetness (Table 1; Table S1) as follows:

a. Rainfall (model 1): Immediate precipitation positively influenced flowering (β = 0.0082, *p* = 0.002), while a 170-day lag showed strong negative effects (β = -0.1324, *p* < 0.0001).
b. Temperature (model 2 and 3): T_max_ had a negative effect on flowering (β = - 1.0636, *p* < 0.0001), whereas T_min_ had a positive effect (β = 1.9474, *p* < 0.001). T_avg_ showed a negative effect (β = -0.9975, *p* = 0.0035), with no lagged associations.
c. Humidity (model 4): Relative humidity had a positive effect on flowering (β = 0.1756, *p* = 0.0076), with no lagged associations.
d. Surface soil Wetness (model 5): Immediate soil moisture strongly promoted flowering (β = 34.722, *p* < 0.0001), contrasting with a negative effect of soil moisture at 20-day lag (β = -21.497, *p* < 0.0001).

**Figure 2:**
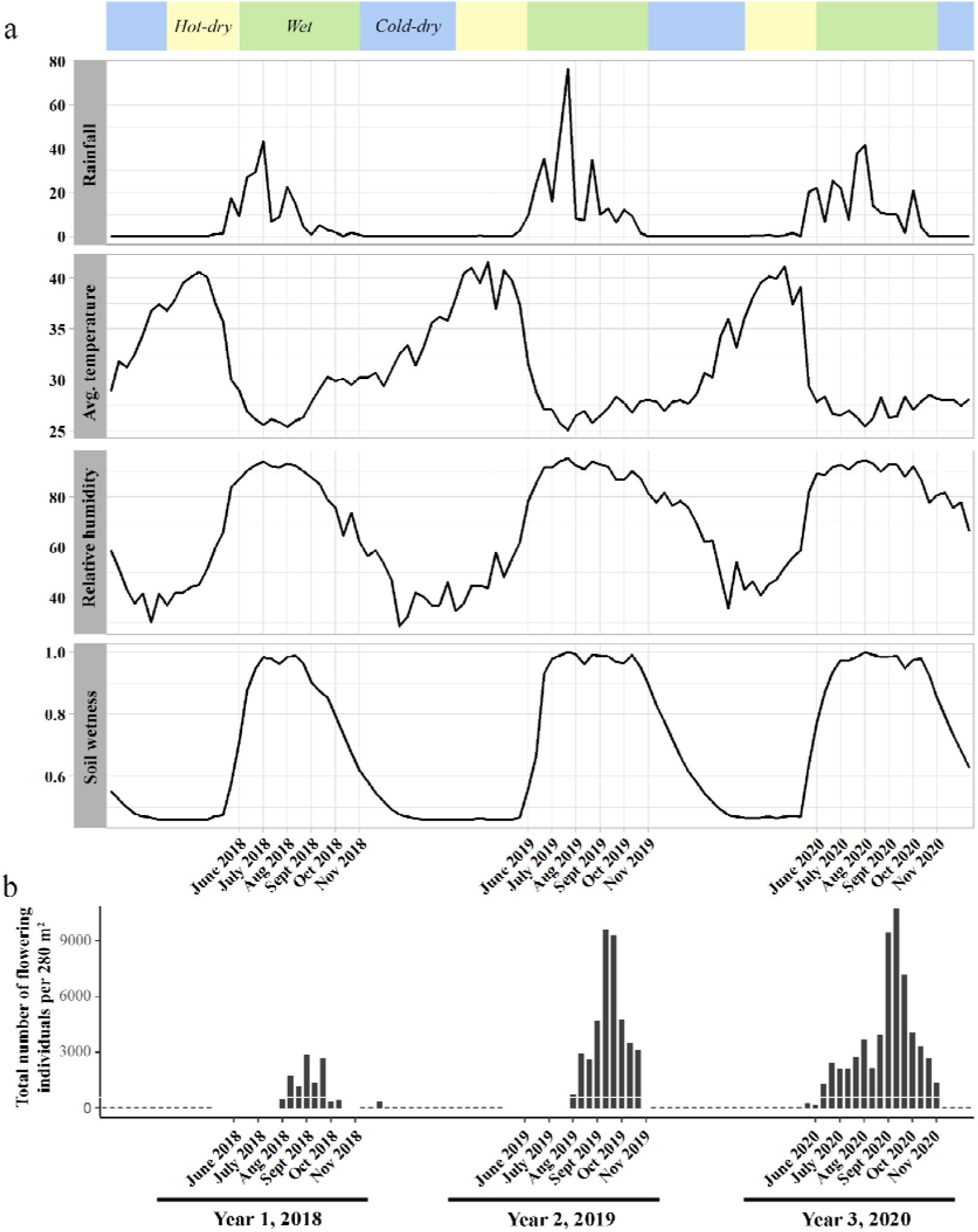
Trends in climatic parameters and flowering phenology across years 2018 to 2020. (a) Daily variations in four climatic parameters - rainfall (mm), average temperature (°C), relative humidity (%), and soil wetness (proportion). Coloured boxes demarcate the seasonal boundaries of wet, hot-dry, and cold-dry seasons. (b) Multi-year trends in total flowering abundance of the study community.

**Table 1:** Parameter estimates and goodness of fit statistics of the GAMLSS (NBI) process for five comparisons to test flowering response to climatic parameters at Kaas. Goodness of fit statistics: GD; global deviance; AIC; Akaike information criterion and SBC; Schwarz Bayesian Criterion. All predictors were significant at *p* ≤ 0.05. Non-significant predictors were excluded and are given in Table S1.

| Predictor | β-estimate | S.E. | *t*-value | *p*-value | GD | AIC | SBC |
| --- | --- | --- | --- | --- | --- | --- | --- |
| **Model 1** |  |  |  |  |  |  |  |
| Intercept | 3.2669 | 0.5529 | 5.909 | <0.0001 | 500.6 | 556.1 | 609.2 |
| **Rainfall** | 0.0082 | 0.0024 | 3.482 | **0.0021** |  |  |  |
| lag17 | -0.1324 | 0.0358 | -3.698 | **0.0012** |  |  |  |
| **Model 2** |  |  |  |  |  |  |  |
| Intercept | 2.856 | 4.1929 | 0.681 | 0.499 | 539 | 581.8 | 629.2 |
| **T<sub>max</sub>** | -1.0636 | 0.1562 | -6.81 | < 0.0001 |  |  |  |
| **T<sub>min</sub>** | 1.9474 | 0.1952 | 9.977 | < 0.0001 |  |  |  |
| **Model 3** |  |  |  |  |  |  |  |
| Intercept | 18.2985 | 4.1134 | 4.448 | < 0.0001 | 571.4 | 600 | 631.4 |
| **T<sub>avg</sub>** | -0.9975 | 0.3261 | -3.059 | **0.0035** |  |  |  |
| **Model 4** |  |  |  |  |  |  |  |
| Intercept | -14.5076 | 2.3417 | -6.195 | **< 0.0001** | 585 | 601 | 618.9 |
| **Relative humidity** | 0.1756 | 0.0636 | 2.763 | **0.0076** |  |  |  |
| **Model 5** |  |  |  |  |  |  |  |
| Intercept | -12.159 | 1.587 | -7.661 | **< 0.0001** | 536.1 | 567.3 | 601.7 |
| **Surface soil wetness** | 34.722 | 3.689 | 9.411 | **< 0.0001** |  |  |  |
| lag2 | -21.497 | 4.354 | -4.938 | **< 0.0001** |  |  |  |

### 3.2. Flowering Phenology

Over three years, we recorded 76 angiosperm taxa on the Kaas plateau, encompassing 57 genera and 31 families (taxonomic details in Table S1). Although 33 taxa showed flowering in all three study years, species richness fluctuated between years, with 53 species in flower in 2018, 43 species in 2019, and 59 species in 2020 (Figure 3; Table S1). The peak flowering abundance also varied across the three years, with the total flowering abundance in 2020 (∼ 10,000 individuals per 280 m^2^) being almost three times higher than in 2018 (∼ 2000 individuals per 280 m^2^; Figure 2b). The only species that consistently displayed high flowering density throughout the season and years was *Eriocaulon sedgwickii* (Eriocaulaceae). Based on the flowering density of each species, we classified 55 species (72.4 %) as nMF (consistently showed low density flowering) and 21 species (27.6 %) as MF species (Figure 3). The number of nMF species recorded each year was 43 species in 2018, 32 species in 2019, and 46 species in 2020 (Figure 3; Table S1). Although *Strobilanthus sessilis* (Acanthaceae) is a well-known supra-annual mass flowering species, we could not detect its mass flowering in our study because the supra-annual frequency of its flowering is about 7 years. The latest recorded episodes for this species on Kaas plateau happened in 2017 and 2024, both of which fall outside of our study period. The species richness of the three colour groups was as follows: Whites (WH = 20 species), Pinks and Purples (PK = 18, PL= 16), and Yellows (YL = 21). While the peak flowering abundance of the community was observed to be in September across the three years (Figure 2b), the peak flowering times of different floral colour groups varied within a year. The WH group showed two flowering peaks in August and in September, the flowering peak of PK and PL was in September, and the flowering peak of YL was in September and October (Figure S4).

**Figure 3:**
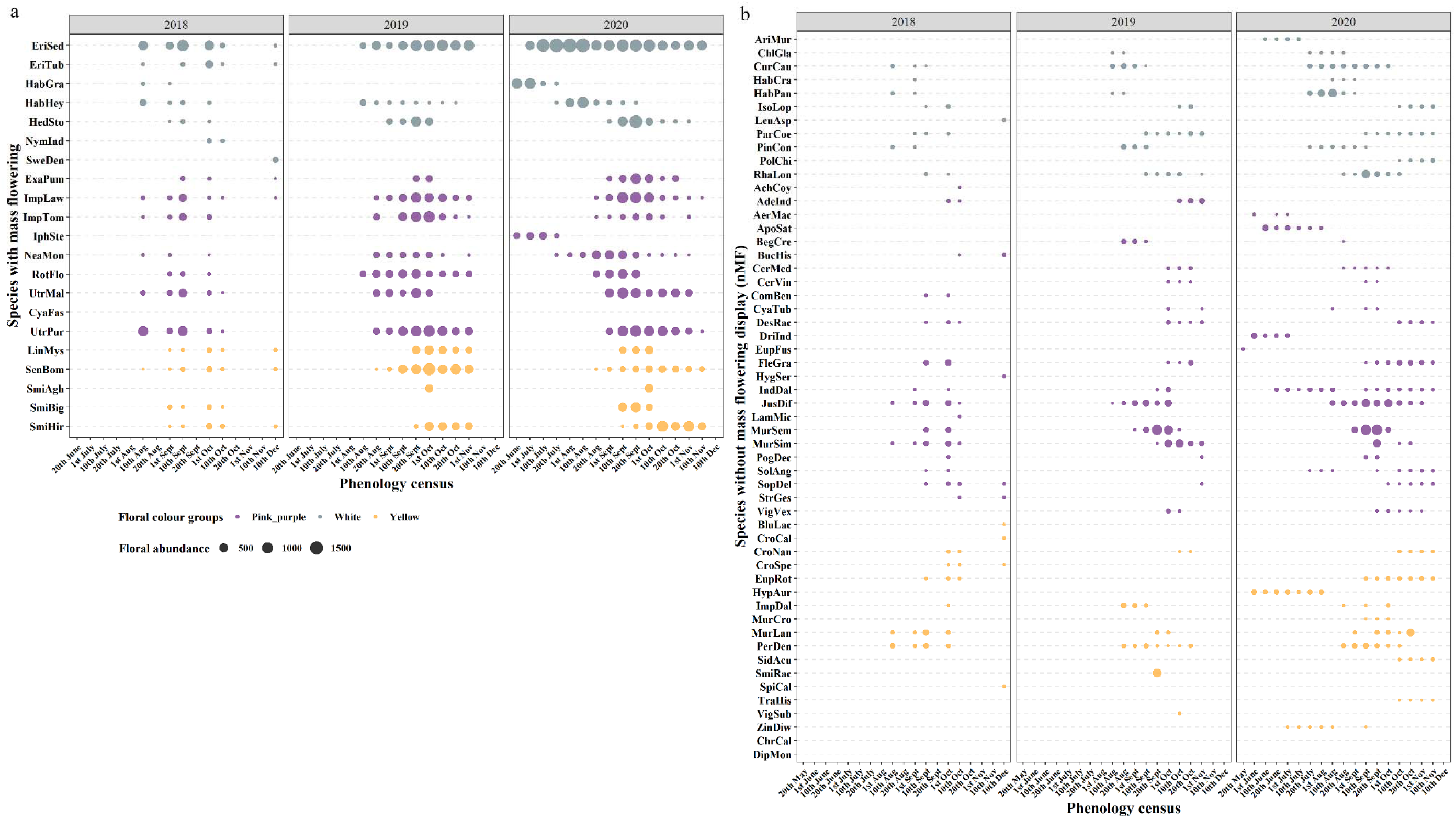
Multi-year trends (2018 to 2020) in flowering phenology of 76 herbaceous species on Kaas plateau, including a) 18 mass flowering (MF) species and b) 58 non-mass flowering (nMF) species. Species names are given as six letter codes and can be referred to Table S2 for their full forms.

### 3.3. Flowering synchrony

We excluded *Eriocaulon sedgwickii* (Eriocaulaceae) from synchrony-analysis due to its consistently high flowering density across years and seasons which resulted in a prominent outlier effect. We found that in all three study years, flowering phenology of the herbaceous community was asynchronous (i.e., SI < 0.4), both in time (SI_temp_ in 2018: 0.3407 ± 0.011; in 2019: 0.3198 ± 0.013; in 2020: 0.2519 ± 0.009) and abundance (SI_abd_ in 2018: 0.147 ± 0.006; in 2019: 0.121 ± 0.007; in 2020: 0.1017 ± 0.005; Figure 4; Table 2). This asynchrony was also found among both MF and nMF species (Figure 4; Table 2). However, MF species displayed significantly higher synchrony than nMF species for both SI_temp_ (Kruskal Wallis chi-squared = 34.9 for 2018, 125.19 for 2019, 43.642 for 2020, d.f. = 1, *p* < 0.05) and SI_abd_ (Kruskal Wallis chi-squared = 46.79 for 2018, 73.882 for 2019, 52.446 for 2020, d.f. = 1, *p* < 0.05) for all three years (Table 2). Synchrony values (SI_temp_ and SI_abd_) when analysed within the three floral colour groups were below 0.4, suggesting an overall low synchrony among species within a color group (Table 2). SI_temp_ values within a colour group was always found to be higher than when compared with the other colour groups, but such trend was not observed for SI_abd_ (Table 2). However, in both of these cases, we did not find any statistical significance.

**Figure 4:**
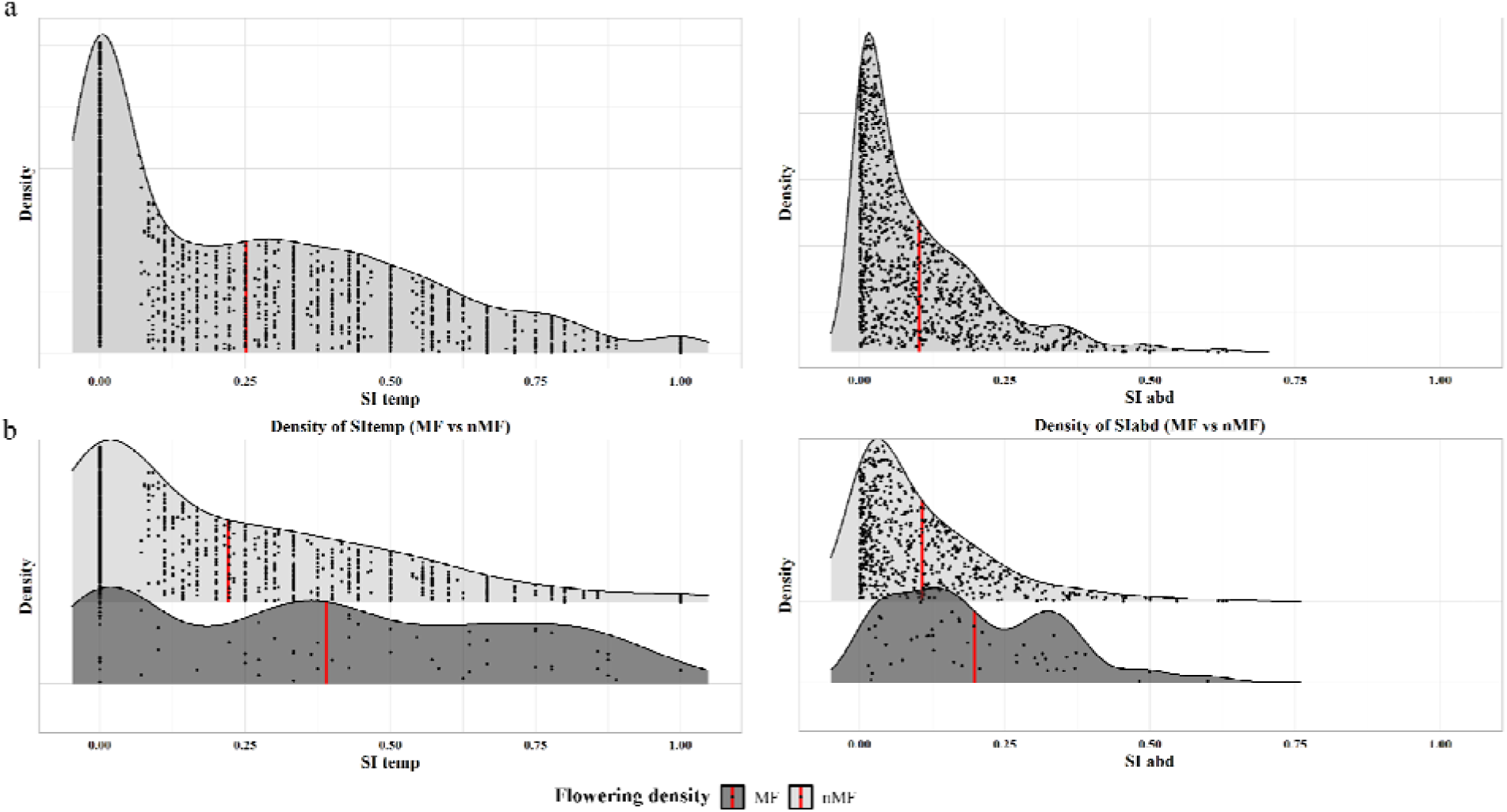
Distribution of SI_temp_ (temporal synchrony) and SI_abd_ (synchrony in abundance) values for: (a) all the species-pairs in the study community, and (b) species-pairs with nMF and MF groups. Mean distribution is shown by the red line. For definition and formulae of the synchrony indices, please refer to Box 1.

**Table 2:** SI_temp_ and SI_abd_ values (mean ± s.e.): (a) the community (all species pairs), (b) MF and nMF pairs, (c) within and (d) between all three floral colour groups.

| **(a) The community and (b) MF and nMF species pairs:** |  |  |  |
| --- | --- | --- | --- |
| **Year** | **Comparisons** | **SI<sub>temp</sub>** | **SI<sub>abd</sub>** |
| **2018** | **Community** | 0.34 ± 0.01 | 0.147 ± 0.01 |
|  | **MF-MF** | 0.562 ± 0.05 | 0.215 ± 0.02 |
|  | **nMF-nMF** | 0.316 ± 0.01 | 0.167 ± 0.01 |
|  | **MF-nMF** | 0.376 ± 0.01 | 0.106 ± 0.004 |
| **2019** | **Community** | $0.32 \pm 0.01$ | $0.121 \pm 0.01$ |
| | **MF-MF** | $0.702 \pm 0.02$ | $0.307 \pm 0.02$ |
| | **nMF-nMF** | $0.248 \pm 0.01$ | $0.125 \pm 0.01$ |
| | **MF-nMF** | $0.377 \pm 0.01$ | $0.088 \pm 0.01$ |
| **2020** | **Community** | $0.25 \pm 0.01$ | $0.102 \pm 0.01$ |
| | **MF-MF** | $0.390 \pm 0.03$ | $0.198 \pm 0.02$ |
| | **nMF-nMF** | $0.221 \pm 0.01$ | $0.107 \pm 0.01$ |
| | **MF-nMF** | $0.293 \pm 0.01$ | $0.081 \pm 0.004$ |
| **(c) Within floral colour groups (intra-group comparisons):** |  |  |  |
| **Floral colour** | **Total number of species** | **SI<sub>temp</sub>** | **SI<sub>abd</sub>** |
| **WH** | 16 | $0.2369 \pm 0.0273$ | $0.1152 \pm 0.0155$ |
| **PK-PL** | 28 | $0.2568 \pm 0.0136$ | $0.0986 \pm 0.0075$ |
| **YL** | 15 | $0.2915 \pm 0.0283$ | $0.0914 \pm 0.0132$ |
| **(d) Between floral colour groups (inter-group comparison):** |  |  |  |
| **Floral colour** | **SI<sub>temp</sub>** | **SI<sub>abd</sub>** |  |
| **WH-YL** | $0.2213 \pm 0.0179$ | $0.1094 \pm 0.0106$ | |
| **WH-PK** | $0.2291 \pm 0.0127$ | $0.1016 \pm 0.0072$ | |
| **YL-PK** | $0.2808 \pm 0.0132$ | $0.1011 \pm 0.0067$ | |

### 3.4. Structure and function of pollination networks

We recorded 90 unique plant-pollinator interactions over approximately ∼ 101 hours of total observation time. Of the 46 flowering species observed, 33 species (71.73 %) received visits from pollinators (Table S3), while 13 species (28.26 %) received no visits. In the cumulative network (Figure S5), we noted a low modularity score (0.365) for the following five modules of interacting pollinators: bee, beetle, fly, ant-wasp, and butterfly-moth. The observed modularity score was significantly lower than those of the randomly generated null model networks (*p* < 0.001). Additionally, null model networks revealed that the observed degree of specialisation (H_2_’ = 0.437; *p* < 0.0001) was significantly lower than expected by chance. At the individual node level (d’), the degree of specialisation ranged from low to moderate, with bees showing the most generalised interactions (d’ = 0.23) and ants exhibiting relatively more specialised interactions (d’ = 0.51; Figure S5; Table S3).

However, among the pollinators, bees (24 links), beetles (20 links), and flies (18 links) were the most frequent visitors (Figure S5). Bees had the highest species strength (10.9), followed by beetles (7.8), flies (7.2), wasps (2.7), ants (2.5), butterflies (1.1), and moths (0.7; Table S3). As the flowering season progressed, the number of species observed increased from 15 in the pre-peak to 25 in the peak season and then decreased to 20 in the post-peak season (Figure 5). A similar trend was observed in the number of links per species (LPS), which increased from pre-peak (1.4) to peak (1.69), and then decreased in the post-peak (1.56) network (Table S3). This shift in plant-pollinator interactions was also evident in the Shannon diversity (H’) values, which increased from 2.9 in the pre-peak to 3.66 in the peak network, followed by a decrease to 3.37 in the post-peak network.

**Figure 5:**
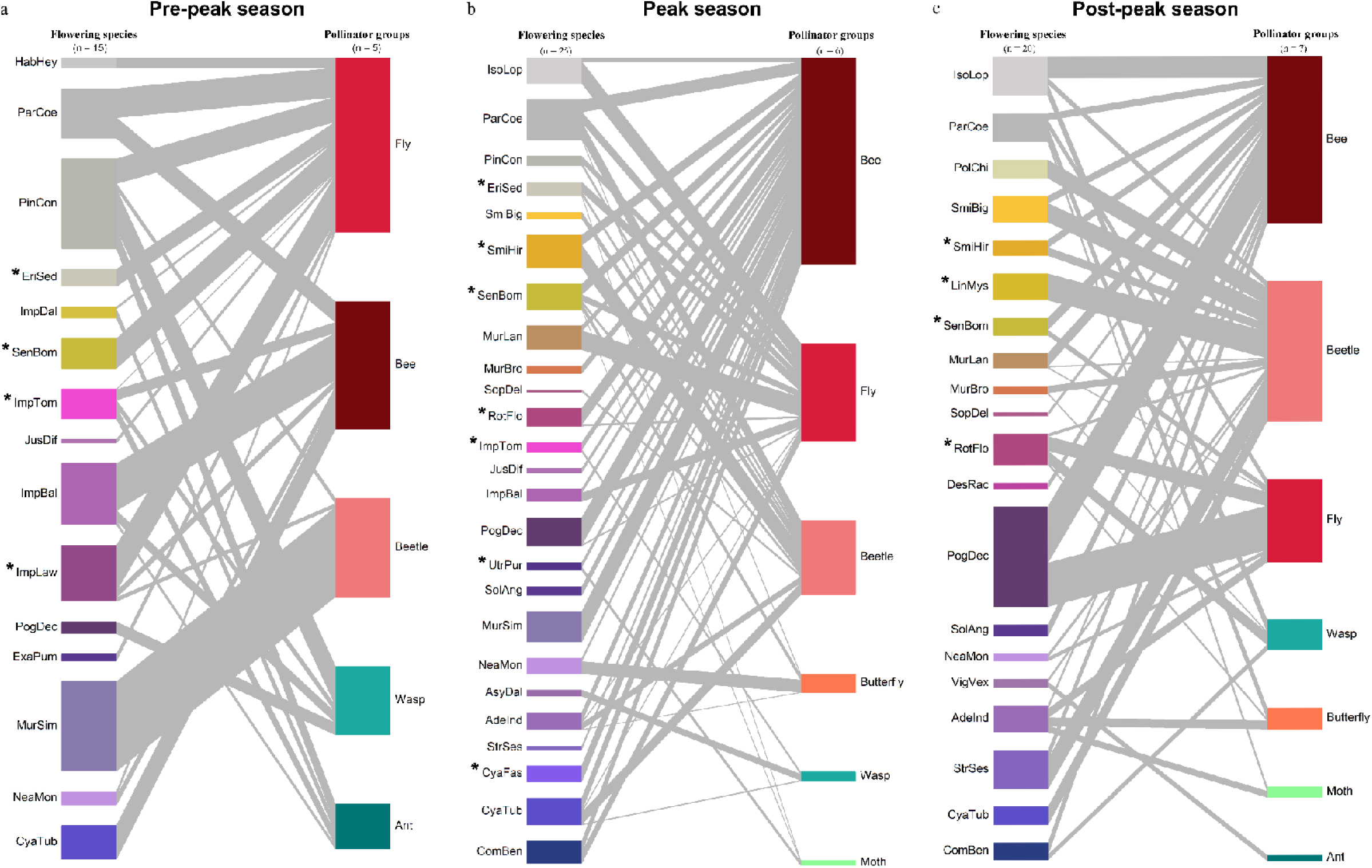
Sub-seasonal plant-pollinator visitation networks for: (a) pre-peak, (b) peak, and (c) post-peak seasons. Total number of nodes in the network are shown by values above the two trophic levels that are flowering species and pollinator groups. The thickness of the links represents the strength of the interaction (here visitation frequency). The colour of the floral colour hue is shown for each plant species, and the size of the coloured rectangles represents the relative importance of the species in the network (that is the number and strength of its interactions with other species). The mass flowering species are denoted by the (*) symbol.

Flies were the most frequent floral visitors in the pre-peak season, with 11 links (Figure 5), while bees became the dominant visitors during the peak (20 links) and post-preak (13 links) seasons. This shift in pollinator relevance was captured by the increasing species strength of bees from pre-peak (3.31) to peak (5.56) and post-peak (7.79) seasons (Table S2). The degree of specialised interactions (H_2_’) was highest in the pre-peak (0.67) network, as compared to the peak (0.56) and post-peak networks (0.57). This trend was supported by higher niche overlap among plants in the peak season (0.43) compared to post-peak (0.40) and pre-peak (0.36), and a greater number of shared pollinators in the peak season (1.02) compared to post-peak (0.92) and pre-peak (0.81; Table S3).

## 4. DISCUSSION

This study presents results from a multi-year monitoring of flowering-phenology at the Kaas plateau where systematic characterization of the unique sky island community was carried out. In addition to flowering phenology studies, the Kaas plateau is also intended as an important long-term monitoring plot for plant-pollinator interactions, trait evolution within the community, and monitoring of adaptive changes within this unique sky-island habitat.

Flowering synchrony between two species is a temporal event, but the ecological effects of this synchrony will also be influenced by the flowering abundance (Marquis, 1988; Elzinga et al., 2007). To emphasise the role of both temporal overlap and abundance values in synchrony, we separated the two factors into separate indices in this study (Box 1) SI_temp_ and SI_abd_. We identified a short flowering phenology on the Kaas plateau that lasted only four to five months, and the community consisted of a *‘few common, many rare’* pattern of species distribution across all years. We report that flowering phenology at Kaas matched yearly shifts in various climatic parameters.

Based on the flowering density of each species we segregated the Kaas plant community into MF species (high flower density) and nMF species (low flower density). We failed to detect any synchrony at the community level, within or between species that show MF or nMF as well as species that share similar floral color. Therefore, competition and asynchrony, and not facilitation by synchrony between similar coloured nMF and MF species seems to be a key phenological characteristic of the plant community on the Kaas plateau.

### 4.1. Seasonality in a Sky Island Community

The short flowering season on Kaas showed a pronounced environmental seasonality, marked by a long dry season (March to May) and subsequent intense Indian monsoon (June to October) and the shifts in flowering phenology also matched yearly shifts in temperature, rainfall, soil wetness, and humidity.

Rainfall has been reported to be critical in the flowering of herbaceous plants in arid and semi-arid landscapes like the tropical Cerrado (Neves et al., 2017), a few tropical seasonal communities (Reich & Borchert, 1984; Hamann, 2004; McLaren & McDonald, 2005; Wright and Calderón, 2006; Kushwaha and Singh, 2008; Stevenson et al., 2008; Lasky et al., 2016; Cortés-Flores et al., 2017), and the desert communities of Northern Oman (Ghazanfar, 1997). Although the Kaas plateau is not perceived as an arid landscape, we propose that the habitat is comparable to arid landscapes due to its strong environmental seasonality and dependence of herbaceous flowering on temperature fluctuations, rainfall, and water availability as well as other geographic features such as rocky landscape with minimal soil layers. In our study, we noted that out of the four climatic factors temperature, rainfall, humidity and soil wetness, three of them are water dependent climatic factors, suggesting the flowering phenology of the Kaas community is majorly water-dependent, alongside temperature.

Between 2018 and 2020, we observed a threefold increase in community-level flowering abundance as well as inter annual variation in flowering density of multiple species. These observations, combined with the noted presence of iconic mass flowering species like *Strobilanthes sessilis* (Acanthaceae) in the community suggest that the Kaas community displays a supra-annual flowering pattern. Similar phenomena have been previously reported in tropical systems (Medway, 1972; Frankie et al., 1974; Ashton et al., 1988; Sakai et al., 1999; Sakai, 2002) and long-term monitoring (>20 years) is critical to confirm these proposed patterns as suggested by Newstrom et al. (1994) and Bawa et al. (2003). Further, while supra-annual flowering has been prominently documented in Southeast Asian dipterocarp forests (Sakai et al., 1999; Sakai, 2002), community-level synchrony in herbaceous-dominated ecosystems of Asia remain understudied, warranting a need for continued phenological studies of the laterite plateaus.

### 4.2. Flora and flowering phenologies within the Kaas community

Species richness of most landscapes report 70% of their species richness to be represented by only 5-6 dominant plant families (Filgueiras, 2002; Royo and Carson, 2005; Oleques et al., 2017). Similarly, on the Kaas plateau, approximately five dominant plant families contributed to 46% of its species richness. The remaining 54% of the species richness was represented by 26 plant families, each of which was further represented by three or fewer species. This pattern also suggests presence of a phylogenetic clustering, which has been noted from high elevational communities (Li et al., 2023). Our future studies will test the presence of phylogenetic clustering among and within the different laterite plateaus of the Western Ghats, which will help in the understanding of the origin and diversification of its flora.

On Kaas, MF species constituted only about 24% of the total species richness (18 species from 11 plant families) while 58 species were nMF species. Since the same species rarely showed mass flowering in consecutive years it suggests that factors such as resource limitation and predator satiation (Janzen, 1976; Augspurger, 1983) that also cause cyclical or periodicity in flowering patterns may play a critical role in the reproductive phenology of the mass flowering species at Kaas. Notably, the ‘super bloom’ like events in Kaas plateau may be a result of individual or a few MF species displaying synchrony in their floral displays. This was coincidentally noted in 2019 when the community showed abundant flowering similar to a super-bloom event which was captured in the high SI_temp_ values recorded for the MF species.

### 4.3. Flowering synchrony

In seasonal communities, climatic constraints can often result in higher interspecific flowering synchrony (Rathcke & Lacey, 1985; Ims, 1990; Vilela et al. 2018; Fisogni et al., 2022). However, despite the strong association observed between environmental seasonality and flowering, a short flowering period, and presence of mass flowering species, our study did not detect interspecific flowering synchrony. Instead, we noted a marked absence of flowering synchrony within the community among the floral color groups and even among MF and nMF species except the year 2019, suggesting a possible absence of pollinator-mediated facilitation through synchrony.

Since flowering asynchrony among species that share floral traits and pollinators can minimise interspecific competition, we suggest that flowering asynchrony in Kaas community may be explained by pollinator-competition hypothesis (Levin and Anderson, 1970; Mosquin, 1971; Stiles, 1977; Anderson and Schelfhout, 1980; Pleasants, 1980; Ashton et al., 1988; Moeller, 2004; Aizen and Rovere, 2010; Pelayo et al., 2021; Arroyo[Correa et al., 2024). Although most studies do not compute synchrony among species, overall flowering asynchrony has been reported in ∼72% of studies across both herbaceous species and woody plants (Feinsinger et al., 1986) and attributed to interspecific competition for pollinators, which remains to be tested in the Kaas community using ecological experiments.

### 4.4. Plant-pollinator interaction

The plant-pollinator networks revealed temporally dynamic interactions that matched the temporal shifts in floral resources within the landscape. We observed higher pollinator sharing when both floral diversity and abundance increased during the peak flowering period, leading to an increase in generalized plant-pollinator interactions. Moreover, the predominance of these generalized interactions highlights the role of asynchrony in minimizing interspecific competition for shared pollinators, which may promote species coexistence in these diverse seasonal habitats. Although generalized plant-pollinator interactions have also been recorded in European grassland communities (Ebeling et al., 2012; Venjakob et al., 2016; Fornoff et al., 2017), studies reporting similar sub-seasonal shifts in interaction networks are limited.

Generalization in plant-pollinator interactions can influence pollen transfer dynamics among species and subsequent reproductive fitness of the plants. For instance, when the same pollinator species visits multiple plant species within a single foraging bout, it can result in higher heterospecific pollen transfers (HPT) and lower fruit set. Although bees formed a dominant pollinator group in Kaas, in a separate study we recorded strong floral constancy among native foraging bees, where individual bees visited the same plant species during foraging bouts (Shrotri et al., 2024), resulting in a possible reduction of HPT rates. We therefore caution that in the absence of empirical data the generalised plant-pollinator interactions detected, should not be interpreted as suggestive of higher HPT rates and lower plant reproductive fitness within this community (Ashman and Arceo-Gomez, 2013; Malecore et al., 2021).

## 5. CONCLUSION

The fragile ecosystems of high-altitude lateritic plateaus in India face immense pressure due to biodiversity loss, habitat destruction, and landscape modification (Watve, 2013). The observed asynchrony in flowering phenology and the absence of facilitative interactions between non-mass flowering and mass flowering species contradicted our predictions that synchrony would be common in these highly seasonal communities. While we detected the community-wide shifts in floral colour which are characteristic of the Kaas plateau, our study shows that this is merely a result of flowering synchrony among some of the MF species (in both SI_temp_ and SI_abd_) and single MF species dominating over the landscape.

Annual plants display an unusual degree of phenotypic plasticity in flowering phenology and this plasticity is adaptive (Schmitt, 1983; Harder and Johnson, 2005; Franks et al., 2007; Blackman, 2017) and serves as a biological indicator of climate change (Gordo and Sanz, 2010; Menzel et al., 2006; 2020). Our study highlights the lack of long-term phenological and ecological studies in the laterite plateaus which restricts its utility for climate change monitoring. Our study is a first step towards characterising these herbaceous communities to understand the ecological and evolutionary mechanisms behind their floristic diversity. Although the genetic basis for mass flowering is not clear, the molecular study in *Strobilanthes flexicaulis* (Acanthaceae) suggests that genetic traits that define periodicity and synchronicity in mass flowering species may be recently acquired from their perennial ancestors (Kakishima et al., 2019). Our current study has now set the stage for future studies to carry out comparative studies that can explore the evolution and maintenance of the herbaceous communities within the Indian subcontinent and across continents.

### Box 1

#### Ambiguity in defining ‘mass flowering’: addressing terminological gaps

The term ‘*mass flowering’* has long been entangled in ambiguity for the following reasons:

1. <u>Common perspective</u>: The definition of the term *mass flowering* is historically rooted in tropical forest studies such as by Ashton et al. (1988) and Appanah (1993). They emphasised *supra-annual flowering*, describing mass flowering as community-wide events tied to multi-year cycles. Similarly, Kelly & Sork (2002) framed masting (and by extension, flowering) as “*synchronous intermittent production*” over decadal scales. Thus, due to its context-specific origins, these definitions capture supra-annual cycles as a *sine qua non* of mass flowering, while perceiving it as a landscape-wide phenomenon without defining the characteristic density traits of the species. This makes the use of the term biologically restrictive by not considering taxa that may show high flowering density across consecutive years in the same habitat, while also neglecting other habitats such as annual herbaceous communities, which are known to form seasonal high density flowering displays.

We believe that Ehrendorfer in Baker (1959) provided a more precise definition of mass flowering as *‘simultaneous flowering of individuals of the same species across a wide landscape where initiation of flowering by certain atmospheric conditions over wide areas.’* Although Enrendorfer’s definition does not convey any information about the flowering duration or periodicity [later addressed by Gentry (1974) and Augspurger (1983)], it unequivocally establishes the density-dependent nature of mass flowering.

2. <u>Contextual bias</u>: Early studies focused on taxa like Dipterocarps or bamboos (Baker, 1959), that show ‘*gregarious flowering*’, which is inherently linked to multi-year cycles. This led to the generalization that *all* mass flowering events must be supra-annual, despite evidence of annual high-density flowering in other systems (Gentry, 1974).

3. <u>Neglect of empirical measures</u>: Classic definitions assumed, but did not measure, high floral density as a hallmark of mass flowering. For example, Ashton et al. (1988) inferred mass flowering from synchronicity rather than abundance, while Baker (1959) misinterpreted “gregarious” flowering of bamboo (a phenomenon exclusively reported in monocarpic plants; Seifriz 1923) with “mass flowering” (a density-dependent phenomenon). Recently, there have been certain attempts to quantitatively estimate ‘masting’ using Coefficient of Variation (CV) and intra-specific flowering synchrony (Herrera et al., 1998; Bogdziewicz et al., 2024). However, in these cases, CV is measured over multiple years for a species by tracking flowering episodes of the same individuals in a given area. Such efforts are possible only for perennial plants and cannot be applied to annual plants or geophytes, where the entire vegetative, flowering, and fruiting phases occur within a short window of 4–5 months or where each individual experiences only one flowering episode per life cycle. Moreover, mass flowering is a dominant phenomenon for seasonal grasslands, for which there is no quantitative measure available currently.

To resolve this ambiguity, we redefine mass flowering as *flowering characterized by empirically measured high floral density*, decoupled from assumptions about periodicity or synchrony. We propose a metric to compute the threshold for classifying a species based on its flowering density (number of individuals in flowering phenophase/m²). A species can be identified as ‘mass flowering’ if its flowering density exceeds 1.64 times the standard deviations above the community’s mean flowering density at a particular time (i.e., z-score of 1.64 per census). This approach is inspired from the remote sensing studies which relies on the threshold to detect outliers or ‘hotspot areas’ using z score (Sims and Thomas, 2002). The threshold effectively identifies species that significantly contribute to larger floral displays compared to other species in the community and can be computed at any given point in the season. Our metric is dynamic and relative to the overall performance of all species in the community; hence it changes with the shifts in flowering phenology. Furthermore, our proposed metric does not distinguish between different life forms and thus can be applied to any system, making cross-system comparisons easier.

##### Distinguishing mass flowering from flowering synchrony

Mass flowering and flowering synchrony represent distinct axes of phenological variation, one relates to quantity, other to timing. Yet they are interchangeably used in literature. In this study we refer to mass flowering as *high floral density* which may or may not coincide with synchrony among species of a community. **Flowering synchrony** refers to *temporal overlap in flowering* (Box 2), which can occur: a) intra-specifically: synchrony among individuals of the same species (a potential contributor to, but not synonymous with, mass flowering) and b) inter-specifically: synchrony across species at the community level—the focus of our study.

Inter-specific synchrony does not imply mass flowering, as co-flowering species may exhibit low density flowering. Conversely, a mass-flowering species may bloom asynchronously relative to others. For example, on the Kaas Plateau, inter-specific synchrony (measured across 2018–2020) remained stable and low due to tight coupling with abiotic cues, whereas flowering density varied annually resulting in shifts in number of species that displayed mass flowering.

### Box 2

#### Methodological gaps in synchrony indices

Existing flowering synchrony indices have critical limitations that restrict their applicability across ecological studies. A prominent issue across multiple methods is the failure to incorporate flowering abundance, as seen in indices proposed by Pool and Rathcke (1979), Primack (1980), Augspurger (1983), and Gorchov (1990), which focus solely on temporal overlap or individual-level metrics. Other indices, such as those by Blomgren (1998) and Freitas & Bolmgren (2008), are constrained to specific scales (e.g., individual or paired comparisons, which is often not applicable to annual herbaceous plants) and lack community-level applicability. Additionally, methods like Mohoro (2002) and Dainou et al. (2012) require high temporal resolution data, limiting their utility for broad-scale or multi-species analyses. Further shortcomings include subjective categorizations of flowering intensity (Michalski and Durka, 2007) and an inability to disentangle within-individual synchrony from population-level patterns (Marquis, 1988).

To address these gaps, we propose two synchrony indices: SI_temp_ (temporal synchrony) and SI_abd_ (synchrony in abundance; Figure B1). SI_temp_ quantifies temporal overlap relative to total flowering duration, resolving issues of skewed phenological distributions (Pool and Rathcke, 1979) and dependence on flowering duration (Augspurger, 1983). SI_abd_ explicitly incorporates flowering abundance by averaging relative abundance ratios only during the co-flowering periods. Both indices are designed for species with high heterogeneity in flowering duration and abundance, decoupling time and abundance to prevent dominance of one factor over the other. Unlike earlier methods, they are applicable across ecological scales (individual, population, community), temporal units (days, months), and community types (herbs, trees), circumventing limitations of taxonomic specificity and restrictive resolution requirements (Mohoro, 2002).

**Figure B1:**
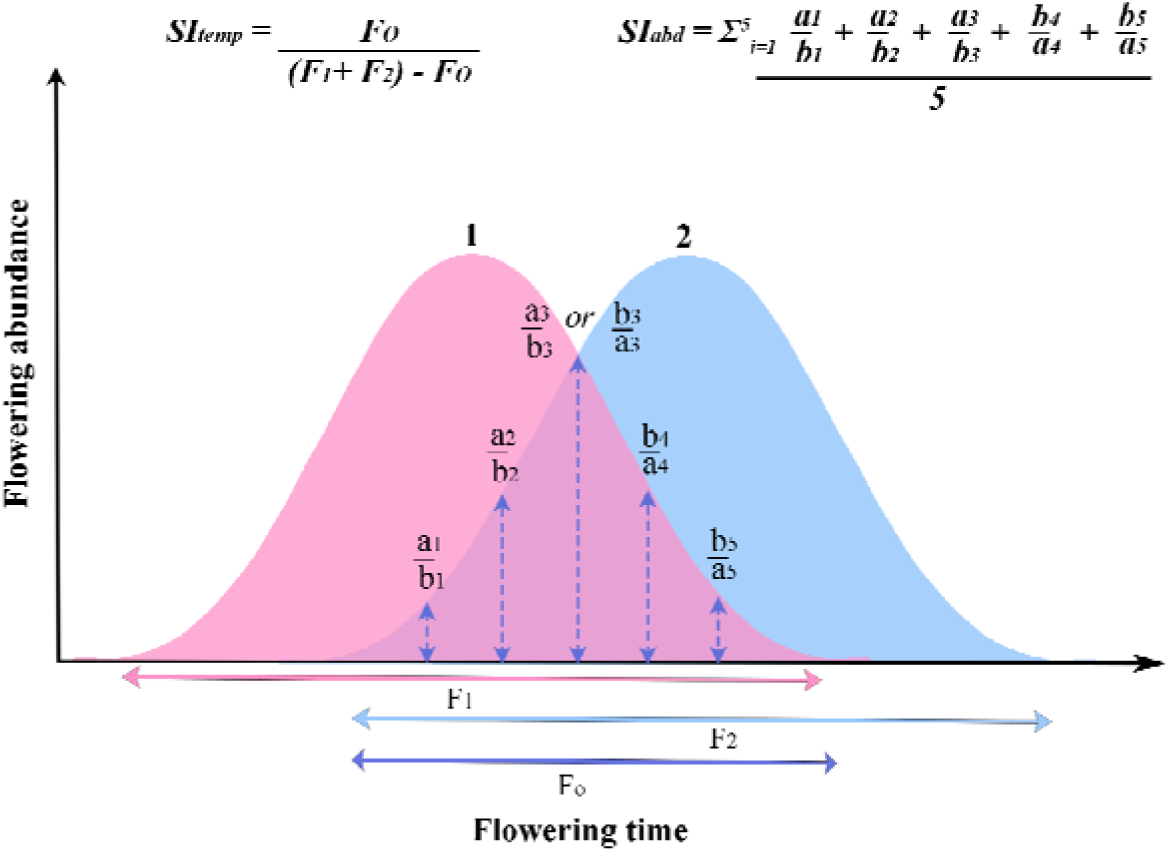
Hypothetical curve of flowering abundance versus time for two species (1 and 2). Species overlap in their flowering time for five censuses, as marked by the dashed lines. *F_1_* and *F_2_* represent the total flowering duration for species 1 and species 2, respectively; and *F_o_* represents the temporal overlap in flowering (i.e., number of days both species are flowering simultaneously).

## ACKNOWLEDGEMENTS

VG thanks the Ministry of Human Resource Development (MHRD) via IISER Bhopal for the start-up grant. VG thanks BD for being an understanding, patient and a personal support throughout this study. SS thanks DST for the INSPIRE fellowship (IF190233) and INLAKS Ravi Sankaran Foundation for INLAKS small grant 2020; SK thanks Council of Scientific and Industrial Research for the Senior Research Fellowship (09/1020(0186)/2019-EMR-1). We thank PCCF, Maharashtra Forest Department, DFO of Satara Division (Forest Department), and the Joint Forest Management Committee (JFMC), Kaas, Satara for research and collection permits [Desk-22(8)/WL/C.R.-16(13-14)/532]. We extend our gratitude to Dr. Hema Somanathan (IISER Thiruvananthapuram) for providing the floral reflectance data for specific species.

## Author Contribution Statement

SS, SK and VG conceived and designed the study. VG and SS acquired the funds and administered the project. RD, NPV, SS and SK conducted the experiments and collected the field data. SS and SK performed the statistical analyses with inputs from VG. SS, SK and VG drafted the manuscript, provided conceptual advice and edited the manuscript. All authors gave final approval for publication and agreed to be held accountable for the contents.

## Open research

All codes used in this study are available via GitHub repository: https://github.com/TrEE-Lab-2025/Phenology_paper_Biotropica. The entire dataset will be available from the authors Gowda, Vinita; Shrotri, Saket; and Kaur, Sukhraj upon request.

## SUPPLEMENTARY INFORMATION

Attached as a separate file; contains three tables, and five figures.

