## Supplementary material for "Mass flowering and flowering asynchrony characterise a seasonal herbaceous community in the Western Ghats": Supp. files

Appendix count: Table: S1 to S3; Figures: S1 to S5.

### Tables

**Table S1:** List of all flowering species along with their species code, plant family, and floral colour groups: Whites (WH), Pinks (PK), Purples (PL), and Yellows (YL). The table indicates the presence or absence of species in phenological transects for the years 2018, 2019, and 2020, along with their mass flowering (MF) and non-mass flowering (nMF) behaviour. The respective year a species displayed MF is marked in bold. If pollinator visitation was observed, it is indicated as Y (yes) or N (no).

| **Sr. No.** | **Flowering species** | **Species code** | **Plant family** | **Floral colour groups** | **2018** | **2019** | **2020** | **Received pollinator visits** |
| --- | --- | --- | --- | --- | --- | --- | --- | --- |
| 1 | *Achyranthes coynei* | AchCoy | Amaranthaceae | PK | nMF | 0 | 0 |  |
| 2 | *Adenoon indicum* | AdeInd | Asteraceae | PL | nMF | nMF | 0 | Y |
| 3 | *Aerides maculosa* | AerMac | Orchidaceae | PK | 0 | 0 | nMF |  |
| 4 | *Aponogeton satarensis* | ApoSat | Aponogetonaceae | PL | 0 | 0 | nMF |  |
| 5 | *Arisaema murrayi* | AriMur | Araceae | WH | 0 | 0 | nMF |  |
| 6 | *Asystasia dalzelliana* | AsyDal | Acanthaceae | PL | 0 | 0 | 0 | Y |
| 7 | *Begonia crenata* | BegCre | Begoniaceae | PK | 0 | nMF | nMF |  |
| 8 | *Blumea lacera* | BluLac | Asteraceae | YL | nMF | 0 | 0 |  |
| 9 | *Buchnera hispida* | BucHis | Orobanchaceae | PL | nMF | 0 | 0 |  |
| 10 | *Ceropegia media* | CerMed | Asclepidaceae | PK | 0 | nMF | nMF | N |
| 11 | *Ceropegia vincaefolia* | CerVin | Asclepidaceae | PK | 0 | nMF | nMF |  |
| 12 | *Chlorophytum glaucoides* | ChlGla | Asparagaceae | WH | 0 | nMF | nMF |  |
| 13 | *Christisonia calcarata* | ChrCal | Orobanchaceae | WH | 0 | 0 | nMF |  |
| 14 | *Commelina benghalensis* | ComBen | Commelinaceae | PL | nMF | 0 | 0 | Y |
| 15 | *Crotalaria calycina* | CroCal | Fabaceae | YL | nMF | 0 | 0 |  |
| 16 | *Crotalaria nana* | CroNan | Fabaceae | YL | nMF | nMF | nMF | N |
| 17 | *Crotalaria spectabilis* | CroSpe | Fabaceae | YL | nMF | 0 | 0 |  |
| 18 | *Curcuma caulina* | CurCau | Zingiberaceae | WH | nMF | nMF | nMF |  |
| 19 | *Cyanotis fasciculata* | CyaFas | Commelinaceae | PL | nMF | **MF** | nMF | Y |
| 20 | *Cyanotis tuberosa* | CyaTub | Commelinaceae | PL | 0 | nMF | nMF | Y |
| 21 | *Desmodium racemosum* | DesRac | Fabaceae | PK | nMF | nMF | nMF | Y |
| 22 | *Dipkadi montanum* | DipMon | Asparagaceae | WH | nMF | nMF | nMF | N |
| 23 | *Drimia indica* | DriInd | Asparagaceae | PK | 0 | 0 | nMF |  |
| 24 | *Drosera indica* | DroInd | Droseraceae | PK | 0 | 0 | nMF |  |
| 25 | *Eriocaulon sedgwickii* | EriSed | Eriocaulonaceae | WH | **MF** | **MF** | **MF** | Y |
| 26 | *Eriocaulon tuberiferum* | EriTub | Eriocaulonaceae | WH | **MF** | 0 | 0 | N |
| 27 | *Euphorbia fusiformis* | EupFus | Euphorbiaceae | PK | 0 | 0 | nMF |  |
| 28 | *Euphorbia rothiana* | EupRot | Euphorbiaceae | YL | nMF | 0 | nMF |  |
| 29 | *Exacum pumilum* | ExaPum | Gentianaceae | PL | nMF | nMF | **MF** | Y |
| 30 | *Flemingia gracilis* | FleGra | Fabaceae | PK | nMF | nMF | nMF |  |
| 31 | *Habenaria crassiflora* | HabCra | Orchidaceae | WH | nMF | 0 | nMF |  |
| 32 | *Habenaria grandifloriformis* | HabGra | Orchidaceae | WH | nMF | 0 | **MF** |  |
| 33 | *Habenaria heyneana* | HabHey | Orchidaceae | WH | nMF | **MF** | **MF** | Y |
| 34 | *Habenaria panchganensis* | HabPan | Orchidaceae | WH | nMF | nMF | nMF |  |
| 35 | *Hedyotis stocksii* | HedSto | Rubiaceae | WH | nMF | **MF** | **MF** | N |
| 36 | *Hygrophila serpyllum* | HygSer | Acanthaceae | PL | nMF | 0 | 0 |  |
| 37 | *Hypoxis aurea* | HypAur | Hypoxidaceae | YL | 0 | 0 | nMF |  |
| 38 | *Impatiens balsamina* | ImpBal | Balsaminaceae | PK | 0 | 0 | 0 | Y |
| 39 | *Impatiens dalzellii* | ImpDal | Balsaminaceae | YL | nMF | nMF | nMF | Y |
| 40 | *Impatiens lawii* | ImpLaw | Balsaminaceae | PK | **MF** | **MF** | **MF** | Y |
| 41 | *Impatiens tomentosa* | ImpTom | Balsaminaceae | PK | nMF | **MF** | nMF | Y |
| 42 | *Indigofera dalzelii* | IndDal | Fabaceae | PK | nMF | nMF | nMF | N |
| 43 | *Iphigenia stellata* | IphSte | Colchicaceae | PL | 0 | 0 | **MF** |  |
| 44 | *Isodon lophanthoides* | IsoLop | Lamiaceae | WH | nMF | nMF | nMF | Y |
| 45 | *Justicia diffusa* | JusDif | Acanthaceae | PK | nMF | nMF | nMF | Y |
| 46 | *Lamprachaenium microephalum* | LamMic | Asteraceae | PK | nMF | 0 | 0 |  |
| 47 | *Leucas aspera* | LeuAsp | Lamiaceae | WH | nMF | 0 | 0 |  |
| 48 | *Linum mysorense* | LinMys | Linaceae | YL | **MF** | nMF | nMF | Y |
| 49 | *Murdannia crocea* | MurCro | Commelinaceae | YL | 0 | 0 | nMF | Y |
| 50 | *Murdannia lanuginosa* | MurLan | Commelinaceae | YL | nMF | nMF | nMF | Y |
| 51 | *Murdannia semiteres* | MurSem | Commelinaceae | PL | nMF | nMF | nMF | N |
| 52 | *Murdannia simplex* | MurSim | Commelinaceae | PL | nMF | nMF | nMF | Y |
| 53 | *Neanotis montholoni* | NeaMon | Rubiaceae | PL | nMF | nMF | **MF** | Y |
| 54 | *Nymphoides indica* | NymInd | Menyanthaceae | WH | **MF** | 0 | 0 |  |
| 55 | *Paracaryopsis coelestina* | ParCoe | Boraginaceae | WH | nMF | nMF | nMF | Y |
| 56 | *Paracaryopsis malabarica* | ParMal | Boraginaceae | PL | 0 | 0 | 0 | N |
| 57 | *Peristylus densus* | PerDen | Orchidaceae | YL | nMF | nMF | nMF | N |
| 58 | *Pinda concanensis* | PinCon | Apiaceae | WH | nMF | nMF | nMF | Y |
| 59 | *Pogostemon deccanensis* | PogDec | Lamiaceae | PL | nMF | nMF | nMF | Y |
| 60 | *Polygonum chinense* | PolChi | Polygonaceae | WH | 0 | 0 | nMF | Y |
| 61 | *Rhamphicarpa longiflora* | RhaLon | Orobanchaceae | WH | nMF | nMF | nMF |  |
| 62 | *Rotala floribunda* | RotFlo | Lythraceae | PK | nMF | **MF** | **MF** | Y |
| 63 | *Senecio bombayensis* | SenBom | Asteraceae | YL | **MF** | **MF** | nMF | Y |
| 64 | *Sida acuta* | SidAcu | Malvaceae | YL | 0 | 0 | nMF |  |
| 65 | *Smithia agharkarii* | SmiAgh | Fabaceae | YL | 0 | nMF | nMF | N |
| 66 | *Smithia bigemina* | SmiBig | Fabaceae | YL | nMF | 0 | **MF** | Y |
| 67 | *Smithia hirsuta* | SmiHir | Fabaceae | YL | **MF** | **MF** | **MF** | Y |
| 68 | *Smithia racemosa* | SmiRac | Fabaceae | YL | 0 | nMF | 0 |  |
| 69 | *Solanum anguvi* | SolAng | Solanaceae | PL | nMF | 0 | nMF | Y |
| 70 | *Sopubia delphiniifolia* | SopDel | Orobanchaceae | PK | nMF | nMF | nMF | Y |
| 71 | *Spilanthes calva* | SpiCal | Asteraceae | YL | nMF | 0 | 0 |  |
| 72 | *Striga gesnerioides* | StrGes | Orobanchaceae | PK | nMF | 0 | 0 | N |
| 73 | *Strobilanthes sessilis* | StrSes | Acanthaceae | PL | 0 | 0 | 0 | Y |
| 74 | *Swertia densifolia* | SweDen | Gentianaceae | WH | **MF** | 0 | 0 |  |
| 75 | *Tragia hispida* | TraHis | Euphorbiaceae | YL | 0 | 0 | nMF |  |
| 76 | *Utricularia malabarica* | UtrMal | Lentibulariaceae | PL | **MF** | **MF** | **MF** | N |
| 77 | *Utricularia purpurascens* | UtrPur | Lentibulariaceae | PL | **MF** | **MF** | **MF** | Y |
| 78 | *Vigna sublobata* | VigSub | Fabaceae | YL | 0 | nMF | 0 | N |
| 79 | *Vigna vexillate* | VigVex | Fabaceae | PL | 0 | nMF | nMF | Y |
| 80 | *Zingiber neesanum* | ZinNee | Zingiberaceae | YL | 0 | 0 | nMF |  |

**Table S2:** Parameter estimates and goodness of fit statistics of the GAMLSS (NBI) process for five comparisons to test flowering response to climatic parameters at Kaas. Goodness of fit statistics: GD; global deviance; AIC; Akaike information criterion and SBC; Schwarz Bayesian Criterion. All predictors that were significant at p ≤ 0.05 are predicted in bold.

| **Predictor** | **β-estimate** | **S.E.** | ***t*-value** | ***p*-value** | **GD** | **AIC** | **SBC** |
| --- | --- | --- | --- | --- | --- | --- | --- |
| **Model 1** |  |  |  |  |  |  |  |
| Intercept | 3.2669 | 0.5529 | 5.909 | **<0.0001** | 500.6 | 556.1 | 609.2 |
| **Rainfall** | 0.0082 | 0.0024 | 3.482 | **0.0021** |  |  |  |
| lag1 | -0.0039 | 0.0026 | -1.49 | 0.1503 |  |  |  |
| lag2 | 0.0009 | 0.0033 | 0.269 | 0.7904 |  |  |  |
| lag3 | 0.0036 | 0.0045 | 0.796 | 0.4345 |  |  |  |
| lag4 | 0.0065 | 0.0056 | 1.163 | 0.2572 |  |  |  |
| lag5 | 0.0017 | 0.0068 | 0.254 | 0.8016 |  |  |  |
| lag6 | 0.0049 | 0.0085 | 0.58 | 0.568 |  |  |  |
| lag7 | -0.0094 | 0.0103 | -0.914 | 0.3708 |  |  |  |
| lag8 | 0.0206 | 0.0119 | 1.725 | 0.0984 |  |  |  |
| lag9 | -0.0134 | 0.0131 | -1.024 | 0.3168 |  |  |  |
| lag10 | 0.0231 | 0.0155 | 1.49 | 0.1502 |  |  |  |
| lag11 | -0.0318 | 0.0159 | -2.002 | 0.0576 |  |  |  |
| lag12 | 0.0269 | 0.015 | 1.787 | 0.0876 |  |  |  |
| lag13 | -0.0329 | 0.0164 | -2.006 | 0.0571 |  |  |  |
| lag14 | 0.0211 | 0.0169 | 1.249 | 0.2246 |  |  |  |
| lag15 | 0.0037 | 0.0176 | 0.211 | 0.835 |  |  |  |
| lag16 | 0.0113 | 0.0219 | 0.519 | 0.6091 |  |  |  |
| lag17 | -0.1324 | 0.0358 | -3.698 | **0.0012** |  |  |  |
| **Model 2** |  |  |  |  |  |  |  |
| Intercept | 2.856 | 4.1929 | 0.681 | 0.499 | 539 | 581.8 | 629.2 |
| **T_max_** | -1.0636 | 0.1562 | -6.81 | **< 0.0001** |  |  |  |
| **T_min_** | 1.9474 | 0.1952 | 9.977 | **< 0.0001** |  |  |  |
| lag1 | -0.0068 | 0.1676 | -0.04 | 0.968 |  |  |  |
| lag2 | -0.0715 | 0.1498 | -0.478 | 0.635 |  |  |  |
| lag3 | -0.0427 | 0.1352 | -0.316 | 0.753 |  |  |  |
| lag4 | -0.0192 | 0.1326 | -0.145 | 0.886 |  |  |  |
| lag5 | -0.1757 | 0.1306 | -1.345 | 0.185 |  |  |  |
| **Model 3** |  |  |  |  |  |  |  |
| Intercept | 18.2985 | 4.1134 | 4.448 | **< 0.0001** | 571.4 | 600 | 631.4 |
| **T_avg_** | -0.9975 | 0.3261 | -3.059 | **0.0035** |  |  |  |
| lag1 | 0.125 | 0.4917 | 0.254 | 0.8004 |  |  |  |
| lag2 | 0.1971 | 0.5111 | 0.386 | 0.7013 |  |  |  |
| lag3 | 0.3643 | 0.4763 | 0.765 | 0.4478 |  |  |  |
| lag4 | 0.1181 | 0.4623 | 0.255 | 0.7994 |  |  |  |
| lag5 | -0.3981 | 0.2873 | -1.385 | 0.1717 |  |  |  |
| **Model 4** |  |  |  |  |  |  |  |
| Intercept | -14.5076 | 2.3417 | -6.195 | **< 0.0001** | 585 | 601 | 618.9 |
| **Relative humidity** | 0.1756 | 0.0636 | 2.763 | **0.0076** |  |  |  |
| lag1 | -0.0214 | 0.0995 | -0.215 | 0.8303 |  |  |  |
| lag2 | 0.1271 | 0.1185 | 1.072 | 0.2879 |  |  |  |
| lag3 | -0.062 | 0.1175 | -0.528 | 0.5997 |  |  |  |
| lag4 | 0.0299 | 0.1379 | 0.217 | 0.8291 |  |  |  |
| lag5 | -0.0001 | 0.0999 | -0.001 | 0.999 |  |  |  |
| **Model 5** |  |  |  |  |  |  |  |
| Intercept | -12.159 | 1.587 | -7.661 | **< 0.0001** | 536.1 | 567.3 | 601.7 |
| **Surface soil wetness** | 34.722 | 3.689 | 9.411 | **< 0.0001** |  |  |  |
| lag1 | -3.987 | 4.572 | -0.872 | 0.3872 |  |  |  |
| lag2 | -21.497 | 4.354 | -4.938 | **< 0.0001** |  |  |  |
| lag3 | 5.604 | 3.512 | 1.596 | 0.1167 |  |  |  |
| lag4 | 1.15 | 3.275 | 0.351 | 0.7268 |  |  |  |
| lag5 | 4.254 | 2.358 | 1.804 | 0.0771 |  |  |  |

**Table S3:** Network indices for plant-pollinator visitation networks—cumulative and sub-seasonal (pre-peak, peak, and post-peak). The indices are calculated at three levels: (a) Network-level, (b) Trophic-level, and (c) Node-level.

| **Indices** | | **Cumulative** | **Pre-peak** | **Peak** | **Post-peak** |
| --- | --- | --- | --- | --- | --- |
| **a. Network-level** | | | | | |
| Links per species (LPS) | | 2.25 | 1.4 | 1.6875 | 1.555556 |
| Modularity | | 0.364542 | 0.558428 | 0.428961 | 0.441068 |
| Shannon diversity (H') | | 4.143325 | 2.901624 | 3.659743 | 3.361387 |
| Degree of specialisation (H_2_') | | 0.437336 | 0.666086 | 0.560439 | 0.567007 |
| **b. Trophic-level (Plants)** | | | | | |
| Number of species | | 33 | 15 | 25 | 20 |
| Mean number of shared partners (pollinators) | | 1.369318 | 0.809524 | 1.023333 | 0.921053 |
| Niche overlap | | 0.407908 | 0.358721 | 0.432176 | 0.401622 |
| **c. Species-level (Pollinator types)** | | | | | |
| Species strength  (number of links) | Ant | 2.494 (7) | 1.515 (4) | 0.818 (3) | 0.667 (1) |
|  | Bee | 10.912 (24) | 3.306 (6) | 12.562 (20) | 7.788 (13) |
|  | Beetle | 7.782 (20) | 1.123 (3) | 4.385 (9) | 6.971 (12) |
|  | Butterfly | 1.133 (5) | 0 | 1.016 (4) | 0.739 (3) |
|  | Fly | 7.184 (19) | 7.222 (11) | 4.443 (12) | 2.306 (7) |
|  | Moth | 0.748 (5) | 0 | 0.482 (3) | 0.379 (2) |
|  | Wasp | 2.746 (10) | 1.835 (4) | 1.293 (3) | 1.151 (4) |
| *d'*  *(node specialisation)* | Ant | 7 | 0.42001 | 0.462261 | 0.862207 |
|  | Bee | 24 | 0.638973 | 0.369795 | 0.380401 |
|  | Beetle | 20 | 0.833492 | 0.52 | 0.718684 |
|  | Butterfly | 5 | 0 | 0.524431 | 0.4275 |
|  | Fly | 19 | 0.615262 | 0.478032 | 0.40918 |
|  | Moth | 5 | 0 | 0.204963 | 0.328391 |
|  | Wasp | 10 | 0.508357 | 0.659442 | 0.452505 |

### Figures

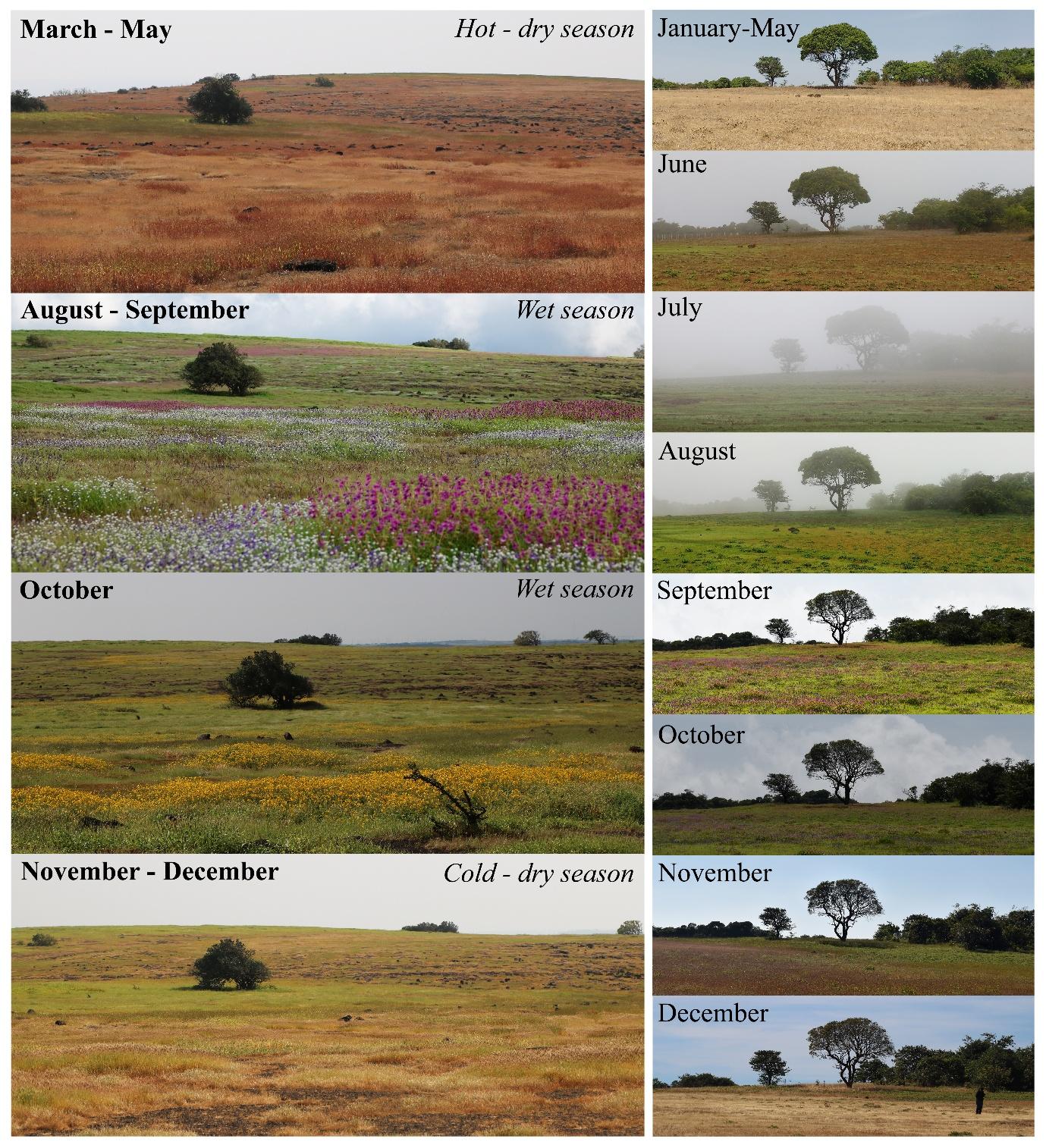

**Figure S1:** Landscape image depicting the shift in vegetation and flowering at the study site—Kaas plateau, over the flowering season (left panel) and within a year (right panel).

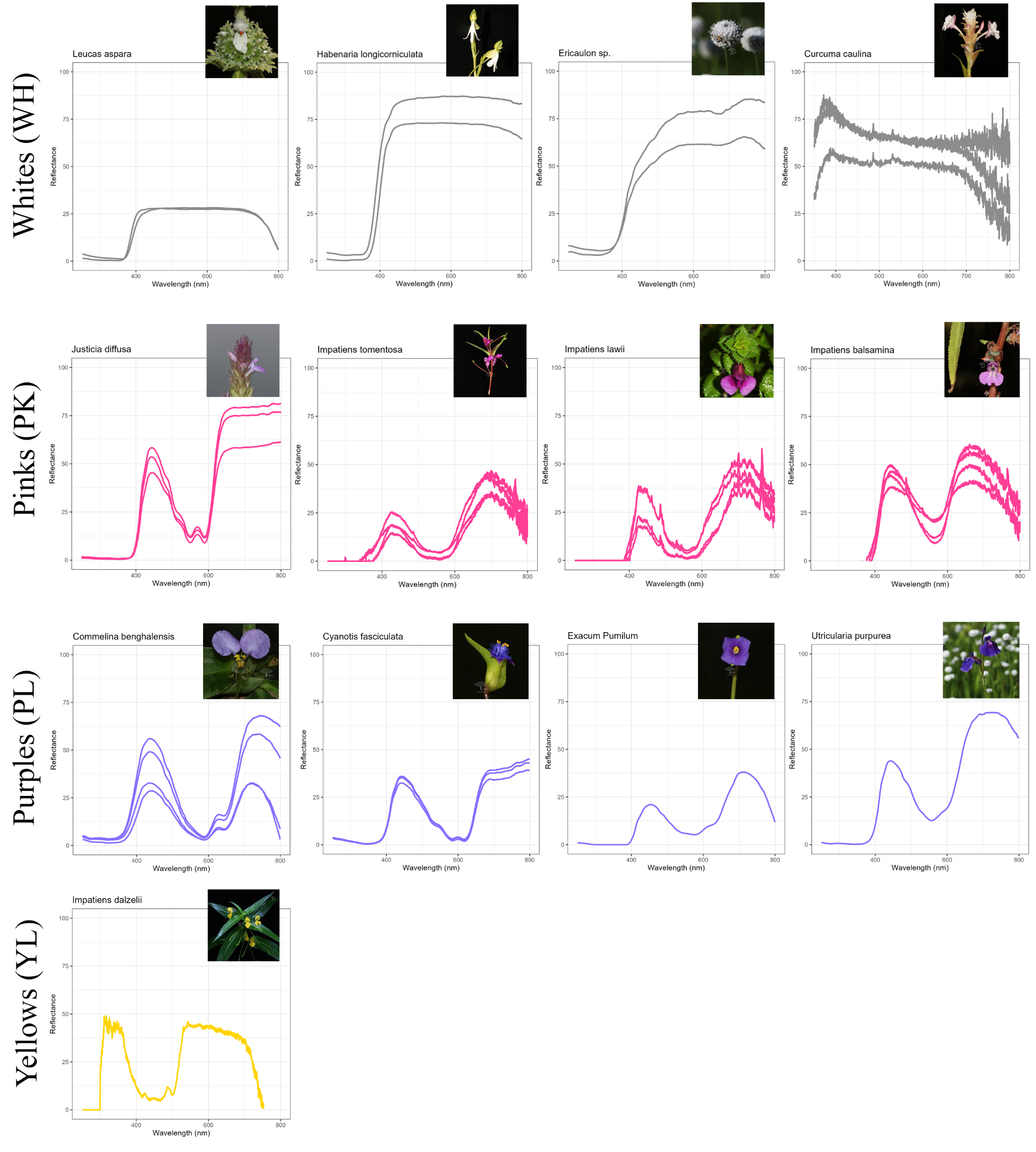

**Figure S2:** Spectral reflectance data for the floral colour groups: Whites (WH), Pinks (PK), Purples (PL), and Yellows (YL).

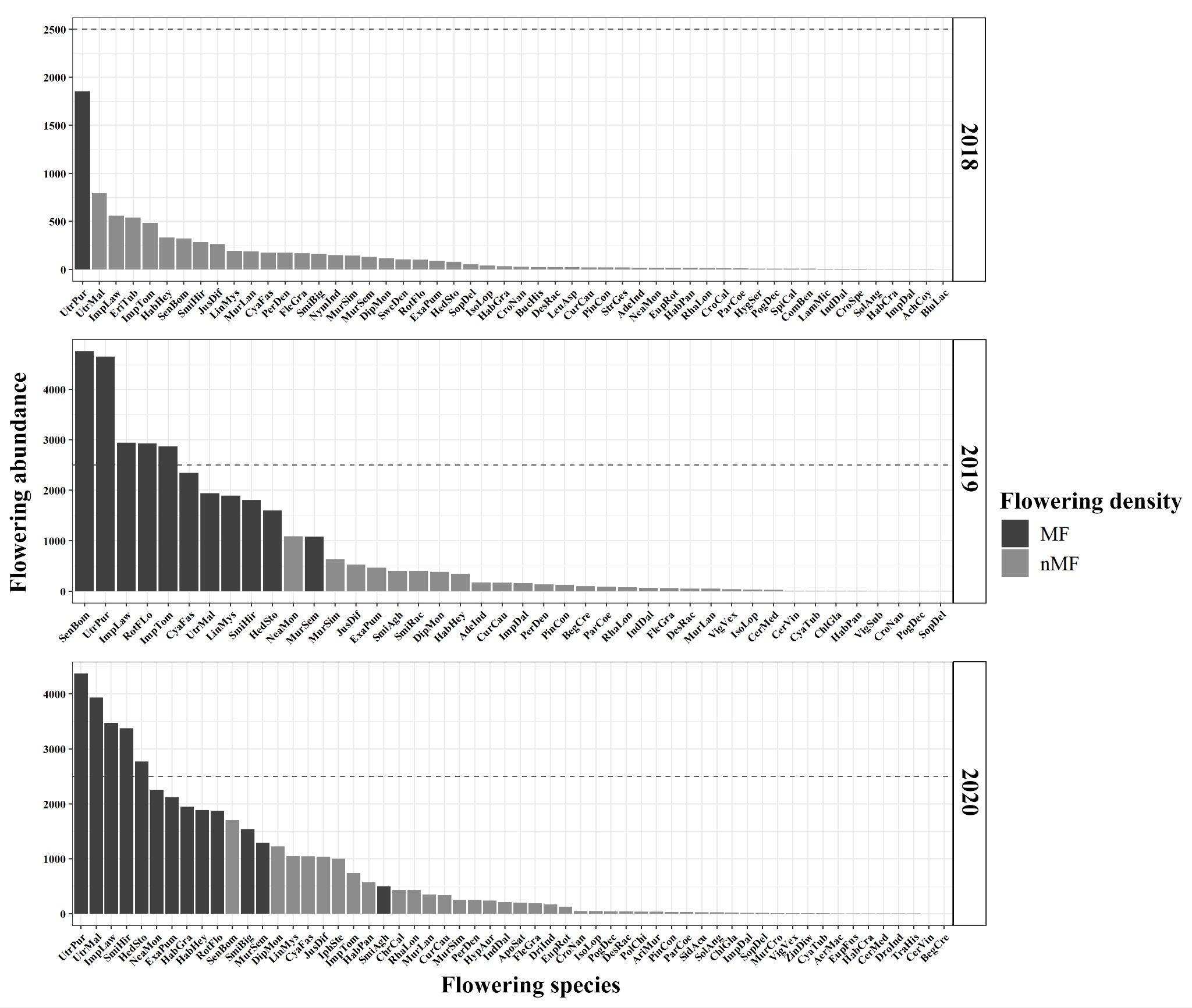

**Figure S3:** Rank-abundance plot displays the *‘few common, many rare’* pattern of species distribution in the community from 2018 to 2020. Dark-shaded bars represent the flowering abundance of mass flowering (MF) species, while light-shaded bars represent non-mass flowering (nMF) species. Dashed lines at a flowering abundance of 2500 serve as a reference point for variation across years.

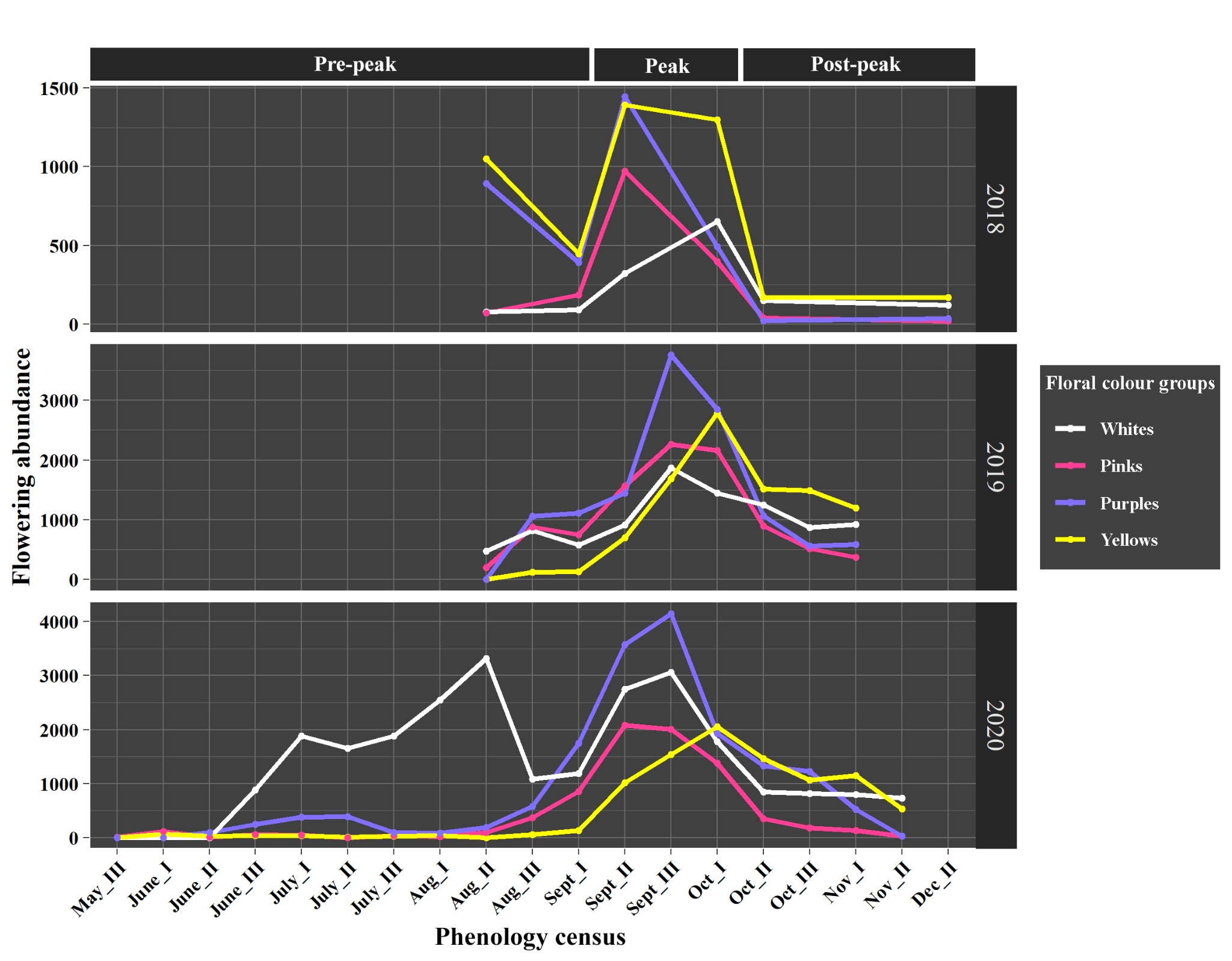

**Figure S4:** Trends in flowering phenology of floral colour groups: whites (WH), pinks-purples (PK-PL), and yellows (YL), over three years from 2018 to 2020.

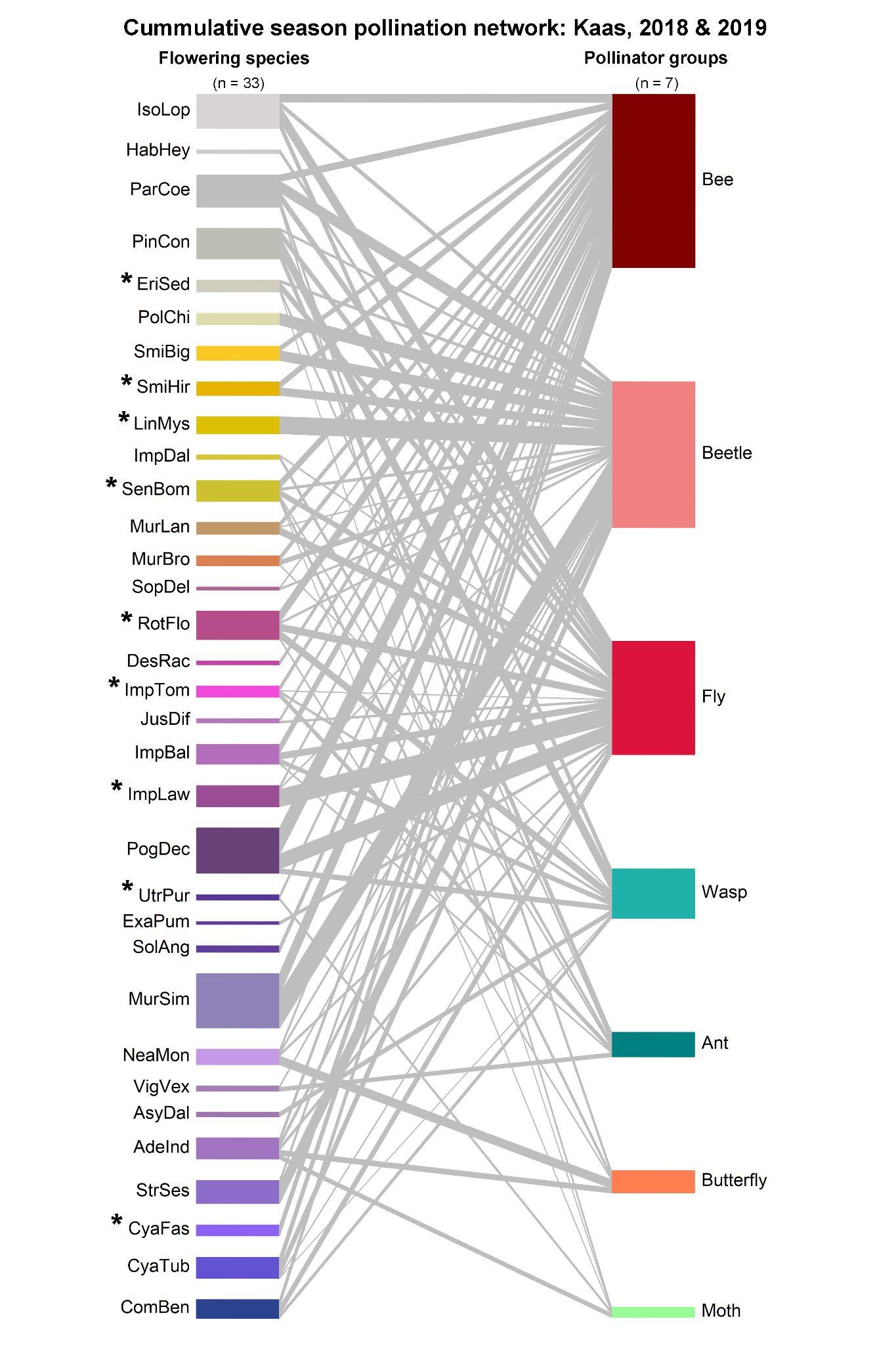

**Figure S5:** Cumulative pollination network for 2018 and 2019. The values mentioned in the bracket above each trophic level denote the total number of species. The MF species are denoted with asterisks (*). The nMF species form 68.89% of the links, while MF species form 31.11%. On assessing the strength of interactions based on pollinator visitation rate, nMF species contributed to 74.29% of the total interaction strength, while MF species represented the remaining 25.91%.
